# Nucleosome placement and polymer mechanics explain genomic contacts on 100kbp scales

**DOI:** 10.1101/2024.09.24.614727

**Authors:** John Corrette, Jiachun Li, Hanjuan Shao, Praveen Krishna Veerasubramanian, Andrew Spakowitz, Timothy L. Downing, Jun Allard

## Abstract

The 3d organization of the genome — in particular, which two regions of DNA are in contact with each other — plays a role in regulating gene expression. Several factors influence genome 3d organization. Nucleosomes (where ∼ 100 basepairs of DNA wrap around histone proteins) bend, twist and compactify chromosomal DNA, altering its polymer mechanics. How much does the positioning of nucleosomes between gene loci influence contacts between those gene loci? And, to what extent are polymer mechanics responsible for this? To address this question, we combine a stochastic polymer mechanics model of chromosomal DNA including twists and wrapping induced by nucleosomes with two data-driven pipelines. The first estimates nucleosome positioning from ATACseq data in regions of high accessibility. Most of the genome is low-accessibility, so we combine this with a novel image analysis method that estimates the distribution of nucleosome spacing from electron microscopy data. There are no fit parameters in the biophysical model. We apply this method to IL6, IL15, CXCL9, and CXCL10, inflammatory marker genes in macrophages, before and after inflammatory stimulation, and compare the predictions with contacts measured by conformation capture experiments (4C-seq). We find that within a 500 kilo-basepairs genomic region, polymer mechanics with nucleosomes can explain 71% of close contacts. These results suggest that, while genome contacts on 100kbp-scales are multifactorial, they may be amenable to mechanistic, physical explanation. Our work also highlights the role of nucleosomes, not just at the loci of interest, but between them, and not just the total number of nucleosomes, but their specific placement. The method generalizes to other genes, and can be used to address whether a contact is under active regulation by the cell (e.g., a macrophage during inflammatory stimulation).

## Introduction

The 3d organization of the genome — in particular, which two regions of DNA are in contact with each other — plays a role in regulating gene expression (1; 2; 3). Many complex, interdependent factors influence genome 3d organization (2; 4; 5; 6; 7; 8; 9; 10), beginning with the polymer mechanics of chromosomal DNA, and including cohesins (in conjunction with DNA binding factors such as CTCF), lamins, phase separation, and nucleosomes, to name just a few. The current understanding of this organization is multi-scale, with nucleosome clutches on the scale of kilobasepairs, topologically associating domains (TADs) on the scale of hundreds of kilobasepairs, compartments on the scale of megabasepairs, and territories on the scale of the genome (11). How DNA contacts are formed and how those contacts support chromatin accessibility through nucleosome remodeling for the binding of regulatory factors remains largely unknown. The full set of biophysical factors that influence genome organization presents a formidable challenge to study. Therefore, to disentangle this complexity, we ask, how much can be explained by nucleosome positioning and polymer mechanics alone?

Nucleosomes are genomic structures comprising ∼ 100 basepairs of DNA wrapped around octameric assemblies of histone proteins (7). While one established role of nucleosomes is to directly control access to a locus (1), nucleosomes also bend, twist and compactify chromosomal DNA, altering its polymer mechanics (7). This raises two possibilities: first, that the number of nucleosomes in between two loci could modulate their contact; second, that specific placement of these nucleosomes, even at approximately the same total number of nucleosomes (i.e., nucleosome frequency), could induce either the formation of new contacts or the loss of existing contacts, as shown in Figure 1. There is mounting evidence for nucleosomes as key contributors in this role, including in physical models (12), synthetic plasmids (13), and experimental perturbations of linker length (9). Compaction by specific nucleosome placement has been shown to encode significant architectural features of the genome in budding yeast (14). These prior studies leave open the question of whether nucleosomes alone can encode long-range genomic contacts in higher eukaryotes.

**Fig. 1:**
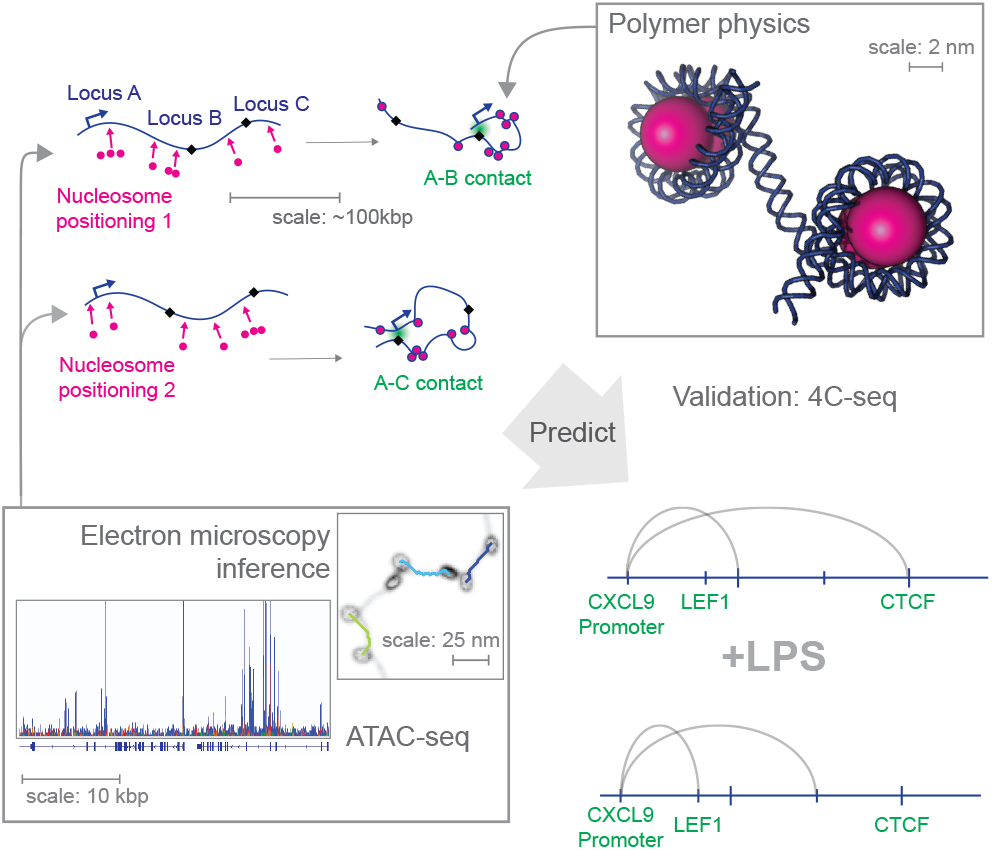
Nucleosomes (pink) between loci introduce “kinks” in chromosomal DNA, potentially driving contacts between distant genomic loci. We find that this is sufficient to significantly modulate the contact of genomic loci on ∼ 100 kilo-basepair scales, and specifically that the positioning of nucleosome, even while holding constant the total number of nucleosomes, can introduce or reduce long-range contacts. Schematic of data driven mechanistic model pipeline: We combine ATAC-seq (24) and electron microscopy (19) data with a mechanistic model of the polymer physics of chromosomal DNA (7) to predict contacts between loci, and validate these predictions with 4C-seq. (Example EM image is synthetic test data.) We find that contact with some transcription factor binding motifs (e.g., LEF1) are gained and some (e.g., CTCF) are lost during inflammatory response in macrophages (“+LPS”).

Furthermore, nucleosome positions change rapidly (within minutes to hours) in response to cellular stimuli, for example during macrophage exposure to pro-inflammatory stimuli (15; 16). This leads us to a third question: to what degree are polymer mechanics and nucleosome positioning predictive of contact after a cell undergoes a functional change (e.g., inflammatory activation in macrophages)? And could these dynamics in nucleosome occupancy coordinate long-range (> 10 kilo-basepair, kbp) contacts between putative regulatory regions associated with active nearby gene promoters?

We address the first possibility by using a stochastic polymer model. Addressing the second possibility requires the polymer model be combined with knowledge of the specific placement of nucleosomes in a particular genomic region of interest. We achieve this with two data-driven pipelines. The first estimates nucleosome positioning from ATAC-seq data in regions of high accessibility (17; 18). Most of the genomic regions are low-accessibility, so we combine this with a novel image analysis pipeline that estimates the distribution of nucleosome spacing from in vivo electron microscopy data (19). With the aid of these two data-driven pipelines, there are no fit parameters in the biophysical model.

Inflammation and the accompanying immune cell activation have provided a useful example for genome organization changes during a cell state transition (20). Herein, we study macrophages undergoing classical pro-inflammatory activation induced by a bacterial cue, lipopolysaccharide (LPS) (21), providing the opportunity to study the transition from unstimulated (before LPS) to stimulated (after LPS) cell states. We apply the method to gene loci for CXCL9, CXCL10, IL6 and IL15, which are inflammatory marker genes that are known to be dynamically regulated in human macrophages following inflammatory stimulation (22; 23). We compare model predictions with contacts measured experimentally by viewpoint-targeted chromatin conformation capture followed by sequencing (4C-seq). We find that within a 500 kilo-basepairs genomic region, polymer mechanics with nucleosome positions can explain 71% of close contacts (*p* = 4 × 10^−8^). Notably, this is not “black box” training (which likely could provide higher predictive power) but rather the consequences only of known physical polymer properties and data-informed nucleosome positions. Therefore, these results suggest that, while genome contacts on 100 kbp scales are multifactorial, they may be amenable to mechanistic, physical explanation.

## Materials and Methods

### Computational simulation of contacts from the twistable worm-like model

We use the twistable worm-like chain model with nucleosomes, as described previously in Beltran et al. (7)^1^. The model is a stochastic polymer model that accounts for twisting and bending mechanics of DNA, as well as the entry and exit angles as DNA interacts with a nucleosome taken from known molecular structure of nucleosomes from crystallography and cryo-EM (25) and mechanical properties of DNA from experiments including optical tweezer experiments (26). This model requires as input the nucleosome positions. Given these, the algorithm uses a chain growth procedure to generate an ensemble of 3d conformations of the DNA between nucleosomes. The configurations between nucleosomes are chosen to obey the thermodynamic canonical ensemble with the energy functional (see Equation 2 in Beltran et al. (7)),

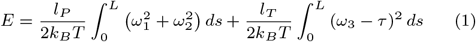

where *l*_*P*_ and *l*_*T*_ are the bending and twisting persistence lengths, *τ* is the twist repeat length, *k*_*B*_*T* is the thermal energy unit, and 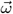 is given by 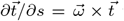 where 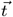 is the tangent vector. The ensemble of configuations can then be queried for statistical quantities like the contact probability between any two loci. Snapshots of this algorithm’s output are shown in Figure 1 (top right), Figure 3I, and Figure 5D.

For the uniform model, we take a mean spacing *µ* and a spacing variability *σ*, and place a nucleosome every *µ* + *X* bp, where *X ∼ U*(*− σ, σ*) and *U* is a uniform random distribution. For the Poisson process, we randomly place nucleosomes every *Y* basepairs where *Y* has a probability distribution *p*_*Y*_ (*y*) = (1*/λ*) exp(*−y/λ*), where our exponential distribution is defined such that the expected value *E*[*Y* ] =*λ*.

Once we randomly sample nucleosomes, we compute the lengths of exposed DNA between nucleosome to generate a list of nucleosome spacings. This list of nucleosome spacings is then provided as input to a chain growth algorithm to compute the 3d conformation of DNA, with nucleosome wrapping, under random thermal fluctuations.

For each conformation, we count each basepair within 10 nm of the locus of interest (usually the TSS). We repeat this a minimum of 150,000 randomly generated polymer conformations and average these scores together. The computation takes approximately 2 hours on an Intel Xeon Gold 6336Y 24-core CPU at 2.4GHz.

### Nucleosome spacing from electron microscopy data

We develop an algorithm for learning nucleosome spacings from EM microscopy data of heterochromatin, specifically here using data from Ou et al. (19). We first determine the centroids of the nucleosomes in the EM data. To do so, we binarize the 3d EM image with a noise threshold of 95% of the maximum voxel value. We find that naked DNA is fully eroded and the nucleosomes cores remain intact in the image. We then use the `cc3d` python package and the function `connected_components` to compute the unique connected 3d components in the image to cluster and compute the centroids of each disjointed candidate nucleosome. With a list of centroids in hand, we return to the original, non-binarized image and use Dijkstra’s algorithm (27) to compute nearest neighbor distances between all centroids. A centroid included in a previous pair is excluded from future pairs. The Dijskstra algorithm weighs all paths by both their intensity under EM and their path-length, and returns the minimal path. We found that this method is the most robust: A method that simply computes shortest paths between detected nucleosomes would ignore DNA curvature; while, on the other hand, many paths are partially disconnected, with gaps that would cause problems with a method that only considers EM intensity.

To validate our method, we simulate DNA in 100 randomly generated 100kbp windows via the ATAC-free algorithm with a uniform nucleosome spacing of 30, 60, and 90 *±* 20 bp (slightly wider distributions than those presented in Figure 2A). We then apply binary dilation with a diameter of 2.5 nm to each 1 cubic nm voxel containing a bp long segment of the polymer, as well as a 5.5 nm dilation to the known positions of the nucleosome cores, so as to match know diameter of DNA and half the diameter of nucleosomes (28; 25). We seek to apply the same noise and uncertainty to the synthetic data as in the real data, which we estimate has a Gaussian blur (point-spead function) width of 1.8nm, and a contrast ratio of 2.5 between nucleosomes and DNA (see Figure S2). We apply Poisson random noise to the synthetic EM image and Gaussian noise ranging from a standard deviation of 0.0 nm to 4.0 nm. We then perform the nucleosome spacing learning algorithm as described above, visually described in Figures S3, S4, and S5. Finally, we compare the resultant learned nucleosome spacing distribution to the simulated ground truth in Figure 3 and Figure S6. At intermediate Gaussian blur levels, the estimate is below the ground truth, due to the construction of our synthetic EM image, where nucleosomes are erroneously connected to themselves. This effect is absent at larger (and smaller) Gaussian blur levels, which more closely represents what the blur that is seen in real ChromEMT data. Overall, these tests demonstrate that the method is able to correctly learn the mean nucleosome spacing for the noise parameters identified from the real data.

**Fig. 2:**
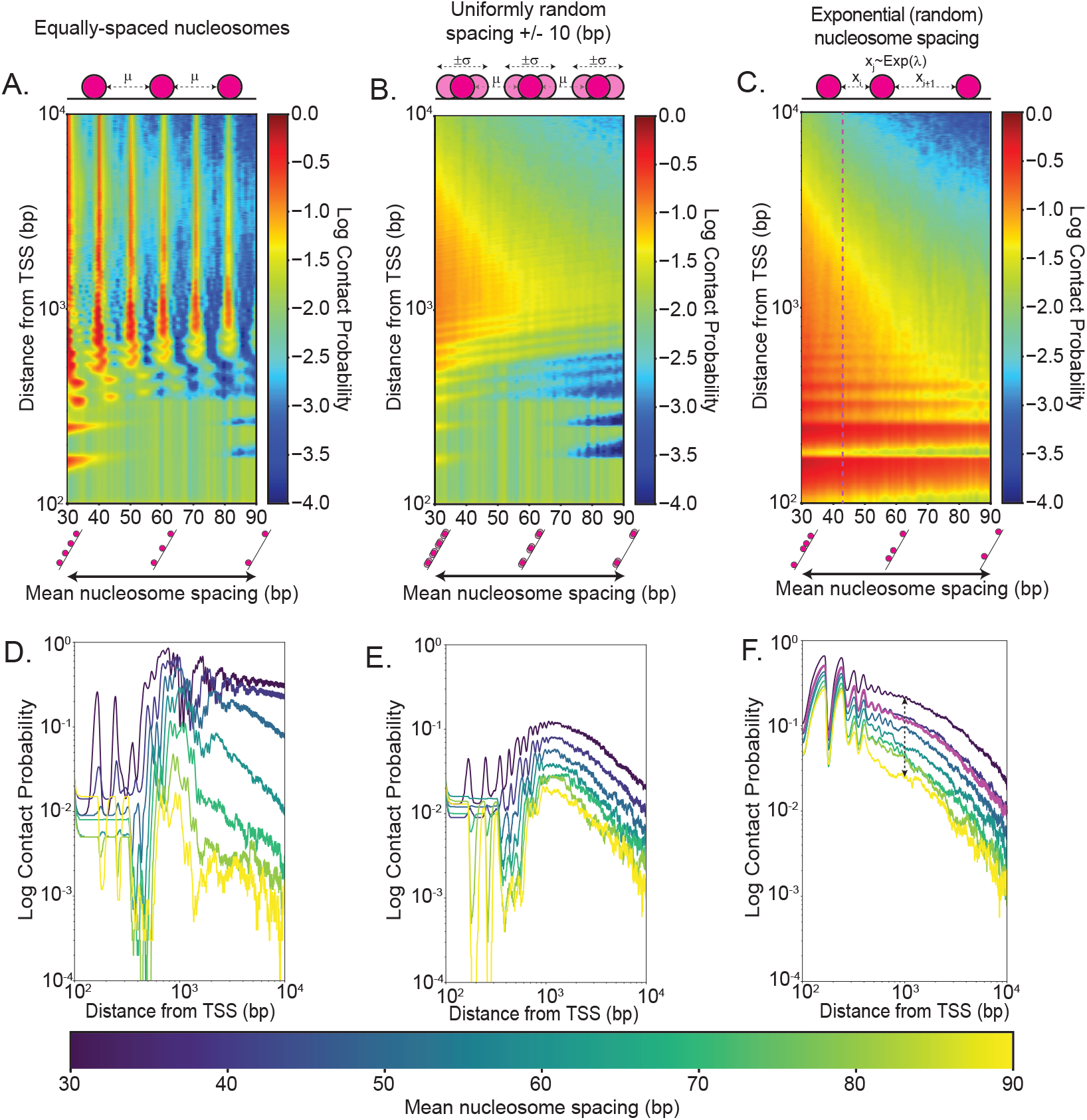
Nucleosome spacing pattern, not just mean spacing, impact long-range DNA contact probability. DNA contact probability of different nucleosome spacing (i.e., linker length) models, defined as the proportion of configurations with the two loci within 10nm, shown as heatmaps (A-C) and families of curves (D-F). **A**,**D** Evenly-spaced nucleosomes with no variation result in a repeated pattern of high contact (red ridges), consistent with the model assumption that the twist repeat of DNA is 10.5 bp. **B**,**E** Introducing variation into the uniform model results in the loss of the spaced ridges, and reveals larger regions of increased and decreased contact probability. **C**,**F** Exponentially-distributed nucleosome spacings (i.e., a spatial Poisson process or lattice gas) result in a smoother contact probability profile. Contact probabilities are highly sensitive to changes in nucleosome spacing, for example changing from 30 bp to 90 bp results in a 88% decrease in contact probability 1kbp away from the locus-of-interest in the Poisson process contact probability model (F, dashed black arrows). The magenta dashed vertical line in C and curve in F indicate the best-fit estimate from our analysis of electron microscopy data (Figure 3).

**Fig. 3:**
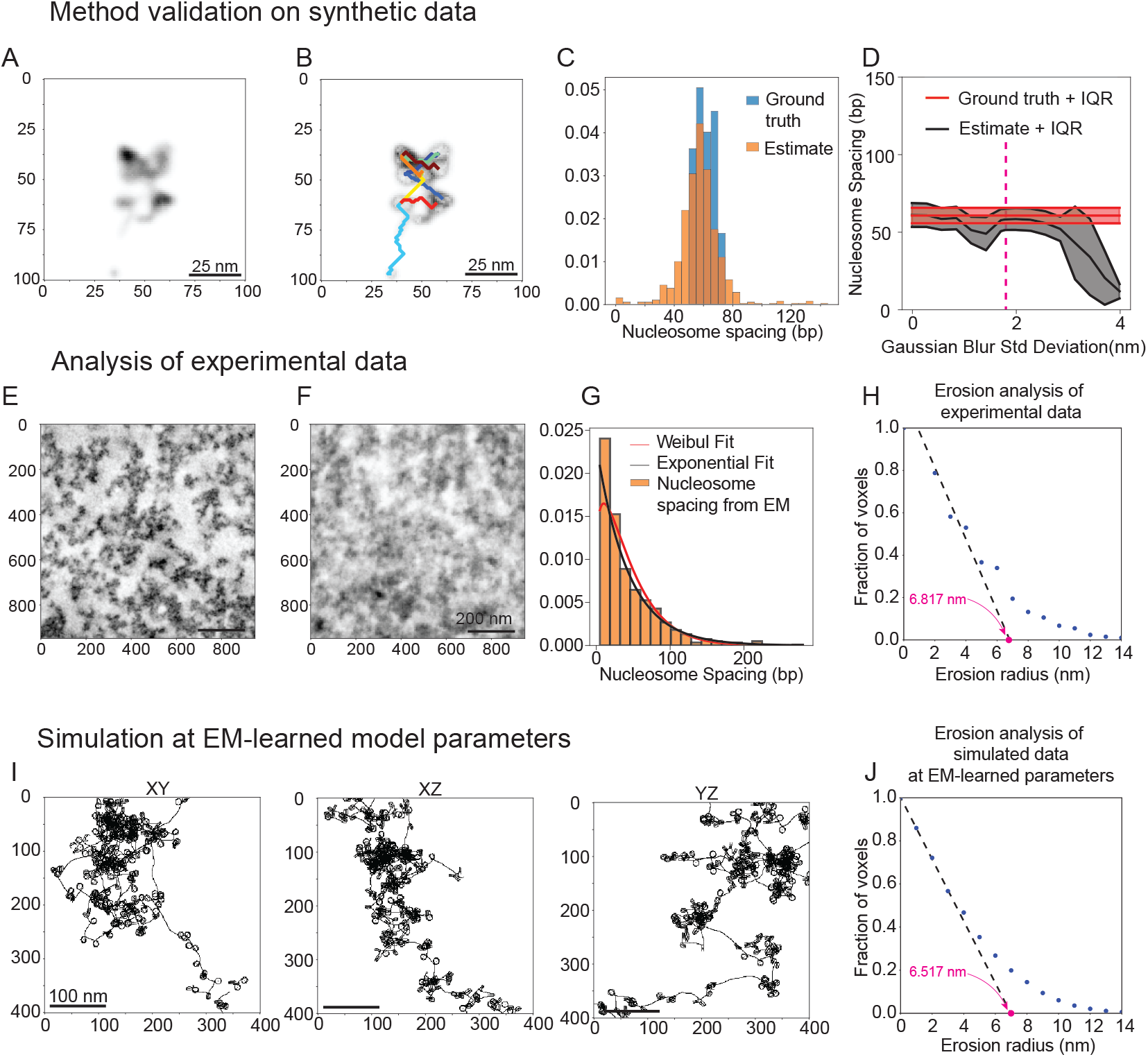
Learning nucleosome spacing distribution from electron microscopy data. **A-D** Validation of image analysis pipeline on synthetic data. **A**. Sample conformation of synthetic DNA with basepair spacing distributed uniform randomly with mean 60 bp *±*20bp, shown as a synthetic EM image with varying voxel intensity. **B**. Sample nucleosome spacing paths estimated by novel Dijkstra’s algorithm pipeline, overlayed on image after estimated nucleosome cores are subtracted (see Methods). **C**. Comparison of ground truth distribution of nucleosome spacing and learned distribution for 100 random synthetic EM images. **D**. The learned nucleosome spacing, summarized by the mean and inter-quartile range (IQR), for various Gaussian blurs. Pink vertical dashed line indicates the point-spread function estimated from the data of Ou et al. (19). **E**. Sample slice of 3d EM microscopy data from Ou et al. (19), **F**. Same as E, after binarization and nucleosome center subtraction for analysis. **G**. Learned distribution of nucleosome spacings. Data fits an exponential distribution with a mean of 43 bp. We also fit a Weibull distribution to the data, where we find our scale parameter to be 1.196, which indicates that our distribution is close to exponential. **H**. Erosion analysis on EM data, following method described by Ou et al. (19). X-axis intercept labeled in magenta (6.817 nm) is indicative of cluster radii in the image. **I**. Sample conformation of synthetic DNA using our learned nucleosome spacing distribution. Data exhibits similar qualitative “clumping” of chromosomal DNA that was identified in Ou et al. (19). **J**. Erosion analysis, same as H, on synthetic data. X-axis intercept labeled in magenta (6.517 nm) is indicative of cluster radii in the synthetic polymers and is within 0.3 nm of the real data in H (6.817 nm).

### Nucleosome Spacing from combined ATAC-seq and EM-informed model

First, we detect peak regions in the ATAC-seq data using MACS2 (29) with a window size of 1kbp. This is to ensure that the regions in ATAC-seq that we determine nucleosome positioning from are of sufficient sequencing depth. We refer to these as high-ATAC-coverage regions. We then use NucleoATAC (17) to determine the probability distribution of nucleosome placement. Briefly, NucleoATAC uses a model fit on *S. cerivisiae* to create estimates of the positions of nucleosomes along the genomic body using ATAC-seq fragment size and locations distributions as input. This model works by running a V-plot, a learned distribution of fragment size relative to the position of a dyad, across the genome, and cross correlates with the genomic spatial fragment size distribution. Thus the output of this model can be interpreted as a probability density function for the locations of nucleosomes across the genome. A higher cross-correlation score coincides with a higher probability with a nucleosome being centered at that genomic position. We verify that our algorithm is behaving as stated by examining the nucleosome densities compared to the NucleoATAC score in Figure S8, as well as verify that the distribution of local maxima in NucleoATAC signal are similar to nucleosome placement probabilities in Figures S9 and S10. Once we have the NucleoATAC probability distribution for a given region of DNA that is high-ATAC-coverage, we sequentially sample the probability distribution for nucleosome center positions until we can no longer place a nucleosome center without being within *±*146 bp (the size of DNA wrapped around the nucleosome) of any previously placed nucleosome center. We then take the remaining regions of DNA that are not high-ATAC-coverage, and fill them with nucleosomes according to a Poisson process (i.e., exponential distribution) with mean 43 bp, as learned from the EM data, as described in the previous section.

### Inflammatory stimulation of THP-1-derived macrophages, 4C-seq library preparation and sequencing

4C-seq was performed according to the protocol described by Krijger et al. (30), with minor modifications. Briefly, 10 million THP-1 cells were seeded in 10 cm dishes and treated with 20 nM phorbol 12-myristate 13-acetate for 48 hours, followed by stimulation with 100 ng/ml ultrapure lipopolysaccharide (Invitrogen, Cat. tlrl-3pelps), which we herein refer to as “stimulated” (versus “unstimulated”). Cells were fixed with 2% paraformaldehyde for 10 minutes at room temperature, quenched with 1 M glycine, and lysed using cell lysis buffer comprising 1M Tris-HCl, NP-40, Triton X-100, 5M NaCl, 0.5M EDTA, and 100X proteinase inhibitor. The supernatant was removed, and 10 ml of cold 0.13 M glycine in PBS was added to quench the reaction, followed by shaking on ice for 10 minutes. Cells were lysed in 5 ml of cold cell lysis buffer, comprising 1M Tris-HCl, NP-40, Triton X-100, 5M NaCl, 0.5M EDTA, and 100X proteinase inhibitor. The mixture was shaken on ice for 15 minutes, then scraped and shaken for another 5 minutes. The nuclei were collected by centrifugation at 500g for 5 minutes at 4^°^C, resuspended in 1.2X restriction enzyme (RE) buffer, and treated with 0.3% SDS and 2.5% Triton X-100, with incubation at 37^°^C between steps.

Chromatin was fragmented by subjecting isolated nuclei to primary restriction enzyme digestion using the 4-bp cutter Csp6I (ThermoFisher, Cat. ER0211) to achieve optimal contact-mapping resolution, followed by in situ ligation in a reaction mix containing ligase buffer and ligase, incubated overnight at 16^°^C. Post-ligation, cross-links were reversed by overnight incubation with Proteinase K at 65^°^C. DNA was purified using SPRI beads and subjected to a second restriction enzyme digestion using NlaIII (NEB, Cat. R0125L) to generate sticky ends for circularization. The digested DNA was then ligated again, and the resulting 3C templates were purified using AMPure XP beads (Beckman Coulter, Cat. A63881), and quantified using the Qubit dsDNA BR Assay Kit. Next, two rounds of PCR amplification were conducted: an inverse PCR to amplify captured regions, followed by an indexing PCR to generate sequencing-ready libraries. Libraries were purified using AMPure XP beads and sequenced on an Illumina MiSeq platform. Data processing, including demultiplexing, trimming, and mapping to the reference genome (hg38), was performed using the pipe4C pipeline as described by Krijger et al. (30). The PCR primers used were:

Primers are shown in Table 1. Universal forward primer for indexing (5’ – 3’): AATGATACGGCGACCACCGAGATCTACACTCTTTCCCTACACGACGCTCTTCCGATCT

**Table 1.**
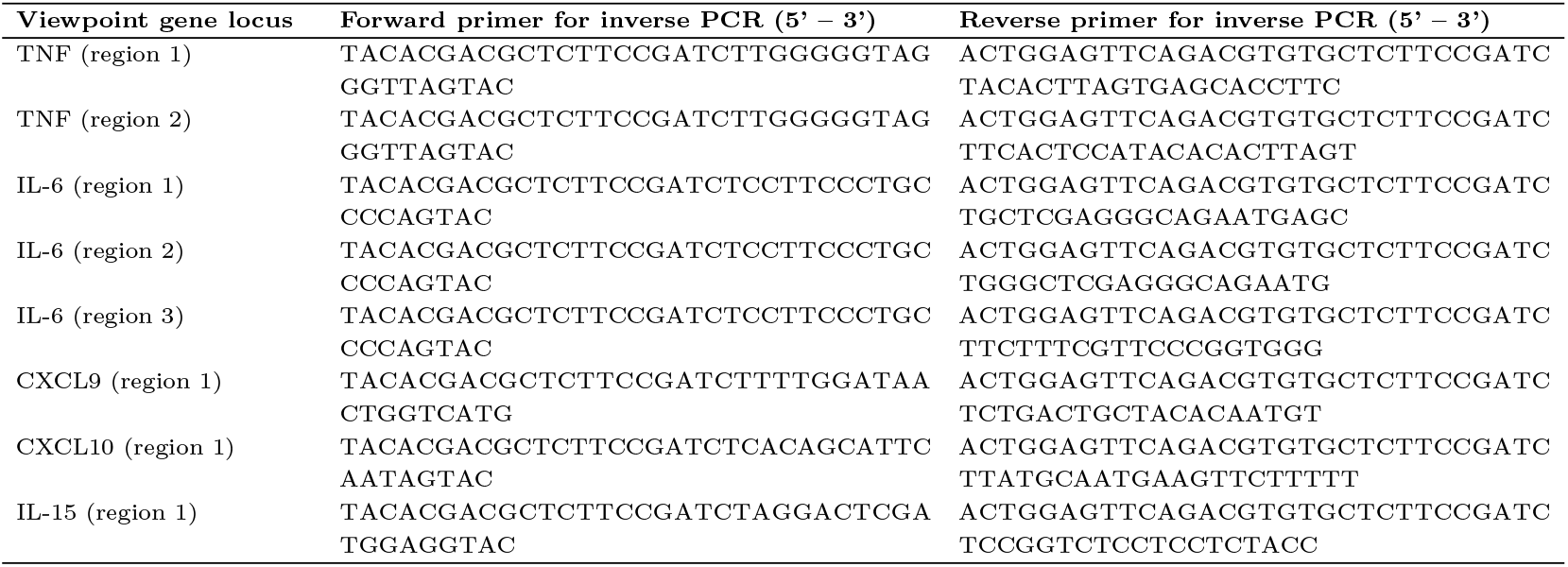
Viewpoint Gene Primers.

### Contact signal processing

To compare our ATAC-informed and experimental 4C-seq contact signals, we seek to identify locations of close contact with the locus of interest. We first perform monotonic regression and subtract the resulting fit from the original data. Monotonic, or isotonic, regression is a technique that fits a piecewise-constant function to data, while enforcing monotonicity. Since we expect contact to peak at the transcriptional start site of each gene and to decay to the left and to the right, we fit two separate monotonic regression models for each side of the site and enforce that they converge to the same peak value at the locus of interest. We do not consider data within *±*10kbp of the locus of interest. We then use a 3rd order Savitzky-Golay filter with a window size of 25kbp. Background removal and smoothing of this nature leads to the data exhibiting heteroscedasticity, with local variance increase further away from the locus of interest (Figure S13B), which can in turn lead to errors in peak finding. Thus, to address this we normalize our data with a rolling z-score with a window size of 25kbp, consistent with our choice of filter window size.

### Contact detection, hyperparameter selection and statistical significance

With data processed as above, use wavelet peak finding with the SciPy function `find_peaks_cwt` (31; 32). This method explores wavelengths in the interval [*α, β*], where *α*,*β* are hyperparameters of the model. We repeat this procedure for both the 4C-seq and the ATAC-informed contact signals.

Although the biophysical model has no fit parameters, the wavelet peak finding algorithm has 2 hyperparameters. To fit hyperparameters *α*, and *β*, we first determine the 4C-seq contact signal parameters by fitting peaks to two replicates of the 4Cseq data, and predicting each other for the 0hr LPS data only. To quantify the predictive performance, for each choice of hyperparameters, we compute the proportion of peaks within 10kbp of replicates one and two respectively, which we term the (ACC) and precision, or positive predictive value (PPV, alternatively 1-FDR), and take the minimum of these as our metric.

In other words, we optimize hyperparameters to be

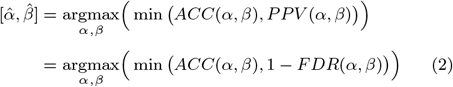

Once we compute 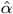 and 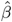 on the 4C data, we use these parameters to compute the peak locations on the average of the 4C LPS 0hr replicates to use as the ground truth.

We then repeat the above optimization where we have the ATAC-informed contact map predicting this 4C-seq ground truth in the same manner.

To estimate the statistical significance of our results, we compare the results of our ATAC-informed model to a null model in which we assume to be uniform random placement of *N* peaks, where *N* is the number of peaks in our estimated 4C ground truth.

We sample *M* different random peak placements, compute the performance metric for each of these, and count the proportion of these placements where this statistic is larger than the performance of the polymer model. In other words, we count the number of random placements with

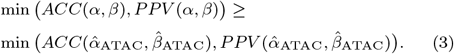

Note that, doing so with the training data (unstimulated cells) is not a valid hypothesis test, since this is the result of an optimization procedure,. We then repeat *p*-value procedure on our test data (LPS-stimulated cells), without refitting our hyperparameters, giving a true statistical *p*-value.

## Results

### Varying nucleosome positioning within the physiologically-observed parameter range leads to order-of-magnitude changes in contact probability

We use an existing model of the polymer mechanics of chromosomal DNA (7) to ask, how much do contacts change when nucleosome positioning is varied within the physiologically-observed parameter range? This model accounts for twisting and bending mechanics of DNA, as well as the entry and exit angles as DNA interacts with a nucleosome, taken from known molecular structure of nucleosomes from crystallography and cryo-EM (25) and mechanical properties of DNA from experiments including optical tweezer experiments (26). This model requires as input the nucleosome positions, while all other parameters are fixed for the purposes of the present work. Given these, the algorithm uses a chain growth procedure to generate an ensemble of 3d conformations of the chromosomal DNA, which can then be queried for statistical quantities like the distribution of distances between any two loci. Snapshots of this algorithm’s output are shown in Figure 1 (top right), Figure 3I, and Figure 5D. See Beltran et al. (7) for full details.

Here, we repurpose this model to look at contact probabilities as a function of distance between two loci. For convenience, we refer to one locus as the locus of interest. Typically, we choose the transcriptional start site (TSS) or a nearby locus. We define the contact probability as the proportion of configurations with the two loci within 10nm. As shown in Supplemental Figure S1, our results are not sensitive to this choice.

Before exploring contact probabilities with nucleosome positions extracted directly from experimental data (next section), here in Figure 2, we perform an *ab initio* study with simplified assumptions for nucleosome positions. In this work, we specify nucleosome positions by specifying the sequence of nucleosome spacings, also called linker lengths, which is the number of basepairs between adjacent nucleosomes, and which does not include 147 basepairs assumed to be occupied by the nucleosome itself. We consider three scenarios: First, we assume nucleosomes are evenly spaced (Figure 2AD). Second, we assume nucleosomes are randomly spaced according to a uniform random distribution with a fixed mean, and a range of *±*10bp (Figure 2BE) following Beltran et al. (7). Finally, we assume nucleosomes are spaced according to an exponential distribution (Figure 2CF). This exponential distribution could arise from a spatially-independent (Poisson) process, or the equilibrium of a 1-dimensional lattice gas. In the simulations where there is no random variation in nucleosome spacing (Figure 2A), contact probability is repeats with a spacing consistent with the model assumption that the twist repeat of DNA is 10.5 bp.

Next, we allow variation where nucleosome spacing is distributed uniformly with a mean *±*10 basepairs (Figure 2B). Here, the local maxima that arose from the twist persistence are “averaged-out”, resulting in a more smoothed-out contact probability profile. We see a local maximum ∼ 1kbp. This maximum occurs when the distance is approximately equal to the persistence length. It coincides with a balance of energetic cost and bending and entropic cost for looping. To elaborate, the persistence length is the length of polymer at which its angle is approximately uniformly random. When the polymer length is approximately equal to the persistence length, it is likely to bend back on itself. If it is much shorter than the persistence length, it sticks too straight to bend over; much longer than its persistence length, it is (on average) too far away in space.

Finally, we allow variation where nucleosome spacing is distributed exponentially (Figure 2C). Some of the qualitative features of the previous nucleosome placement models are preserved, but with a larger region of increased contact around the locus of interest. The maximum observed in Figure 2A and 2B is absent. As more randomness is added to the polymer mechanics (in this case, via nucleosomes), the persistence length becomes more random, so the critical length that maximizes looping becomes random, and no maximum is observed in the heatmap.

Taken together, these results demonstrate that contact probabilities are highly sensitive to changes in nucleosome spacing. For example, changing from 30 bp to 90 bp results in a decrease in contact probability 1kbp away from the TSS of 88%, almost ten-fold (Figure 2F, dashed black arrows). These results also demonstrate the important role of nucleosome spacing pattern, and not just mean spacing or density. Motivated by these, we next seek to estimate the spacing distrubtion in cells.

### Analysis of existing electron microscopy images allows learning nucleosome spacing in low-ATAC-coverage regions

To put the previous results into the context of a specific gene, in a cell undergoing regulated changes in gene expression, we require a method to extract nucleosome positions from data. This is possible using the DNA accessibility sequencing method ATAC-seq (24) and algorithms such as NucleoATAC (17) or NucPosSimulator (18). However, we found that most of the genomic landscape (e.g. *≈* 98% for within *±*200kbp of IL6 TSS) has insufficient ATAC-seq data coverage, defined as any region that of the ATAC-seq data that is not identified as a peak region by MACS2 peak calling (29). Therefore, for these low-ATAC-coverage regions, we develop a nucleosome spacing learning algorithm that uses electron microscopy (EM) images of chromosomal DNA, combined with existing EM images (19).

The full method is described in the Methods. Briefly, we first binarize this data to a threshold such that only the dark core of the nucleosome remains, and perform 3d connected components analysis to identify each distinct candidate nucleosome. We then apply a novel iterative pathfinding algorithm on the original, unbinarized image. This algorithm weighs all paths by both their path-length and their intensity under EM (where darker, i.e., denser regions are weighted less), and returns the minimal path between candidate nucleosomes identified from the binarized image. This proved advantageous: A method that simply computes shortest paths between detected nucleosomes would ignore DNA curvature, while a method that only considers EM intensity fails on many nucleosome pairs where the path is disjoint (due to imaging noise).

We test the method on synthetic data with a known ground-truth distribution of nucleosome spacings (Figure 3A-D), finding that the learned distribution is similar to the ground truth in terms of its mean and interquartile range, with larger tails in the learned distribution (Figure 3C,D). Testing on synthetic data is further elaborated in Methods and Figures S2-S6.

We then apply the method to EM images from Ou et al. (19), taken from heterochromatin in U2OS cells. We use a region of 328 *×* 328 *×* 119.72 nm, resulting in *≈* 1400 nucleosomes (Figure 3E). We find a nucleosome spacing distribution that is well-approximated as exponential, with a mean of *≈* 43 bp, as shown in Figure 3G and indicated in Figure 2C,F. This learned distribution of exponential with mean of 43 bp is in the range of values predicted and/or assumed in previous work (33; 34; 8).

We simulated the polymer mechanics model at the learned distribution and parameters, and then we use this to conduct comparisons with the original data. Besides the qualitative comparison in Figure 3I showing similar clutch sizes, we provided two quantitative comparisons.

First, we repeat the erosion analysis in Ou et al. (19) on both the real data (Figure 3H) and synthetic data (Figure 3J). Briefly, in erosion analysis, the binarized 3d image is convolved with a sphere of increasing radius, and volume of the new image (which has been eroded) is measured. The resulting volume as a function of erosion radius is then fitted with a linear function for the first five data points as done in Ou et al. (19). We specifically quantify the comparison with the horizontal axis intercepts, finding radius 6.8 nm and 6.5 nm in the real data and synthetic polymer models, respectively.

Second, we compute the chromatin packing scaling factor *D* as defined in Li et al. (35). This scaling factor estimates the fractal behavior of a polymer. Previously, Li et al. (35) determined that *D ≈* 2.6 before a transition around autocorrelation radius of 102.4 nm. As Li et al. (35) point out, the packing factor cannot be directly measured in simulated polymers, but it can be approximated as the inverse of the scaling exponent of the end-to-end-distance. Using their described methods, we observe a similar transition length, as shown in Figure S7, and, using this, we estimate *D ≈* 2.8.

### An ATAC/EM-informed polymer model to understand the effects of nucleosome positioning and polymer mechanics on long-range contact probabilities and loci of high-contact for specific genes

The above method allows the study of nucleosome-driven contacts for a specific gene. We first apply the method to CXCL9, a gene that is upregulated in response to inflammatory stimulation (22; 23). This gene, along with the other genes we study, were chosen for their known role in inflammation as well as sufficient ATAC-seq reads for nucleosome position estimation (16). The full pipeline for simulation uses ATAC-seq data to place nucleosomes on regions of high-ATAC-coverage with the NucleoATAC algorithm (17), and then uses the EM-informed model presented above for regions of low-ATAC-coverage, resulting in a complete map of nucleosome placements around the CXCL9 TSS. Our algorithm for sampling nucleosome placements from NucleoATAC in regions of high-ATAC-coverage is described in Methods. In Supplemental Figures S8-S10 we show that our nucleosome placement algorithm is consistent with the NucleoATAC data. We can then perform the same stochastic polymer mechanics calculation as in Figure 2 but with data-informed nucleosome positions. Using published ATAC-seq data from Zhang et al. (16), we perform this simulation for ATAC-seq data from both unstimulated macrophages, and from macrophages stimulated with bacterial lipopolysaccharide (LPS) for 1 hour. Crucially, there are no fit parameters in the biophysical model, since DNA mechanical properties are fixed by previous experiments (25; 26; 7).

Note that, while Mnase-seq may provide an alternative and more direct measurement of nucleosome positions, a method that uses ATAC-seq is valuable, since ATAC-seq data is more prominent (by a rough estimate, the GEO database has around 17,000 Mnase-seq datasets and 64,000 ATAC-seq datasets).

The configurations, as viewed by radius of gyration and root-mean-square distances, show discrepancy from the 0.5 power-law predicted by the simplistic Gaussian chain assumption (Supplemental Figure S11), consistent with simulations of chromosomal DNA in budding yeast (14). We plot the data-informed contact probabilities in Figure 4C. For comparison, we overplot the contact probabilities predicted for the exponentially-distributed nucleosome spacing distribution estimated from EM data alone (so, same as Figure 2F), which approximately obeys Gaussian chain statistics. The discrepancies between these two nucleosome placement assumptions demonstrates that real nucleosome placement can indeed significantly modulate contact, both quantitatively and in introducing new qualitative features, e.g., a sudden reduction in contact around -25kbp.

**Fig. 4:**
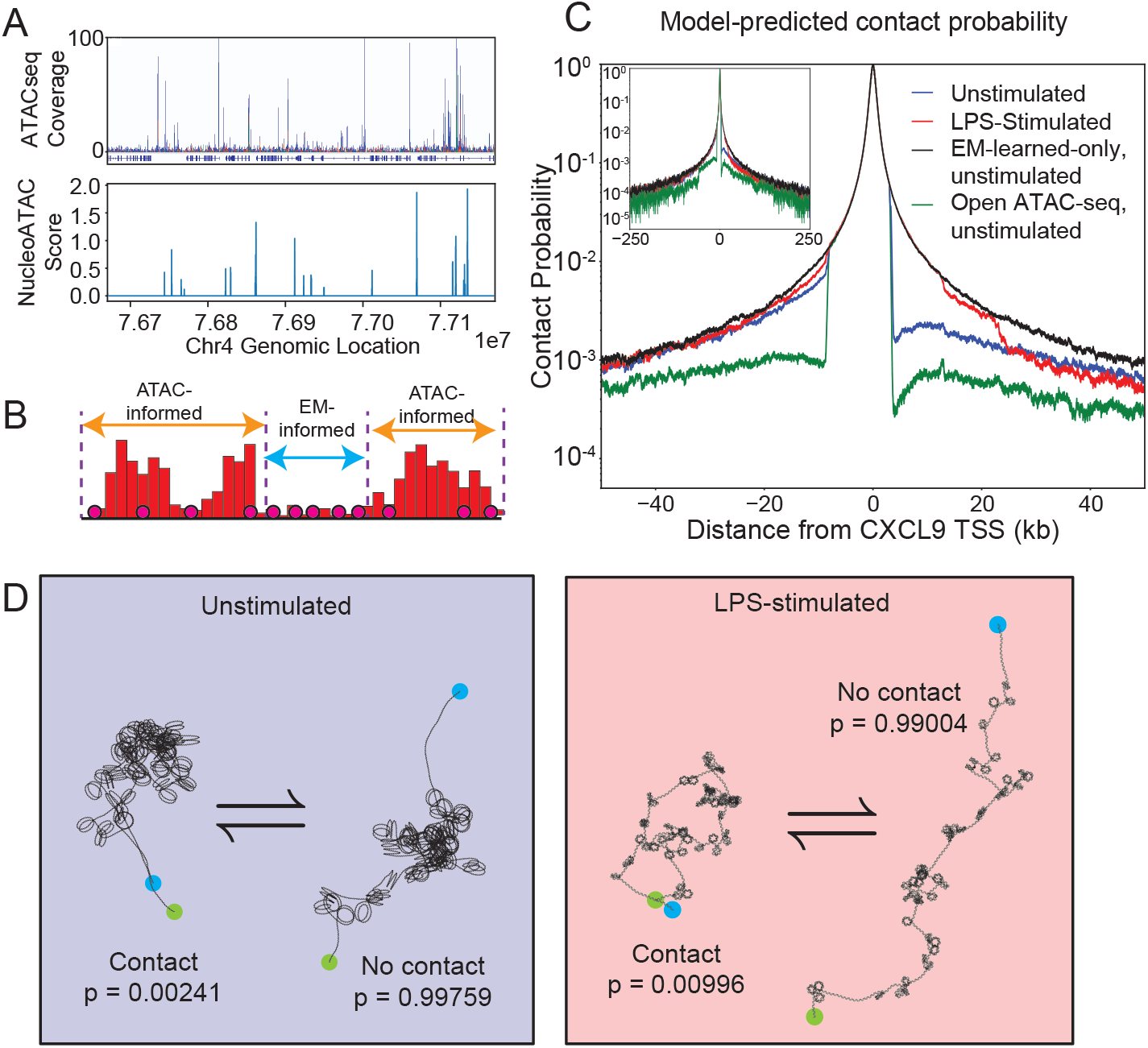
Simulation of polymer mechanics of CXCL9 with nucleosomes placed according to ATAC-seq and EM data. **A**. Example ATAC-seq coverage (top) and NucleoATAC output probability density function (bottom) centered at CXCL9 TSS *±*250kbp. Peaks correspond with a higher probability of nucleosome presence in transcriptionally active DNA. ATAC-seq and NucleoATAC maps are sparse and require EM data to fill in low coverage regions. **B**. The MACS2-identified high-coverage regions are analyzed using NucleoATAC (17), which gives a probability distribution of nucleosome center positions in all peak regions (orange). We sample these regions for nucleosomes, then fill the remaining EM-informed regions (blue) with nucleosomes according to the learned distribution (described in Figure 3). This list of nucleosome spacings for the genomic region of interest of DNA is provided to the polymer mechanics model (7), and we perform Monte Carlo sampling of the 3d conformation of DNA in thermal equilibrium. **C**. Contact probability profiles for unstimulated (pre-LPS; blue) and 1-hour post-LPS stimulated macrophages (red) within a *±*50kbp window around the transcriptional start site of CXCL9. EM-only (black) and open ATAC-seq (green) is shown for comparison. Open ATAC-seq corresponds to not placing any nucleosomes in MACS2-identified high-coverage regions as seen in B. The EM-only probability profile approximately behaves as a Gaussian chain. Inset is the same data but plotted on a *±*250kbp window to demonstrate the genomic domain of simulation. **D**. Sample conformations of CXCL9 before and after LPS-stimulation. One example locus is shown in contact with the transcriptional start site; this contact probability quadruples upon LPS-stimulation, suggesting that nucleosome placement alone is sufficient to significantly alter contact probabilities.

The raw contact probabilities show significant differences between the contact probabilities before and after LPS stimulation (blue and red curves in Figure 4C). One example locus is shown in contact with the transcriptional start site in Figure 4D. This contact probability quadruples upon LPS-stimulation, suggesting that nucleosome placement alone is sufficient to significantly alter contact probabilities. Note that this is not simply a consequence of increasing or decreasing the number of nucleosomes, since the total number of nucleosomes varies less than 2% between the two conditions.

To further study differences in contacts before and after LPS stimulation, we produce Hi-C-like contact maps in order to study the overall conformational changes our model predicts (Supplemental Figure S12). In these contact maps, we see that on average the polymers form associated domains with edges coinciding with locations of NucleoATAC peaks. In other words, our polymer morphology is primarily determined by the accessible regions of DNA input into our model via ATAC-seq. We note that when compared to other works with Hi-C, Micro-C, or other methods, such as Goel et al. (36), our predicted contact maps decay faster, and the associated domain-like structures are not as strongly defined. We attribute this to our model not accounting for spacial restrictions on the polymers, such as a nuclear envelope, as well as no nucleosome-nucleosome interactions being explicitly modeled.

To further characterize the conformations, we simulate 2d (i.e., pairwise) contact maps before and after stimulation in Supplemental Figure S12. We observe association domains, which appear as triangles along the bottom edge of Supplemental Figure S12. These association domains coincide with locations of NucleoATAC signal, representing regions of reduced nucleosome occupancy.

We can compare the simulation with Hi-C, Micro-C or other high-throughput conformation capture methods, such as data in Goel et al. (36), with several caveats. First, the domain size and resolution are different, with Hi-C on larger domains than the 500kbp domain simulated here. Second, we must compare relative changes in contact signal, since absolute contact signal depends on molecular details of the methods. Third, in both our work (Figure 4C) and others (36), there is significant heterogeneity across genomic regions. With those caveats, the predicted contact maps in our nucleosome-only simulation decay faster (from 125kbp to 250kbp, roughly 5-fold reduction in contact probability) compared to, e.g., Figure 2B in Goel et al. (36) (from 125kbp to 250kbp, roughly 2-fold reduction in contact signal). This implies that our simulated conformations are more expanded, suggesting that spatial restrictions not included in our model play a role. (Surprisingly, as presented below, this expanded conformation provides an intermediate step in accurate close-contact location prediction.)

### The ATAC/EM-informed polymer model is predictive of long range genomic contacts observed by 4C-seq

We next sought to identify specific loci that we could categorize as being in “close contact” with the locus of interest, namely the CXCL9 transcriptional start site. We perform background removal, smoothing, and normalization to produce a contact signal (see Methods and Supplemental Figure S13). We then use wavelet peak finding (32) to identify “peaks” in the contact signal, which we then refer to as predicted close contacts.

Note that while the biophysical model that predicted the contact probabilities shown in Figure 4C has no fit parameters, there are 2 hyperparameters in the wavelet peak finding algorithm. To avoid overfitting, we optimize these hyperparameters on 4C data before LPS stimulation (Figure 5ACE, “Unstimulated”, which serves as the training data set of LPS-treated human macrophages at zero hours), and then use the same peak-finding hyperparameters on one hour post-LPS-treated cells (Figure 5BDF, “Stimulated”, which therefore serves as the validation data set).

**Fig. 5:**
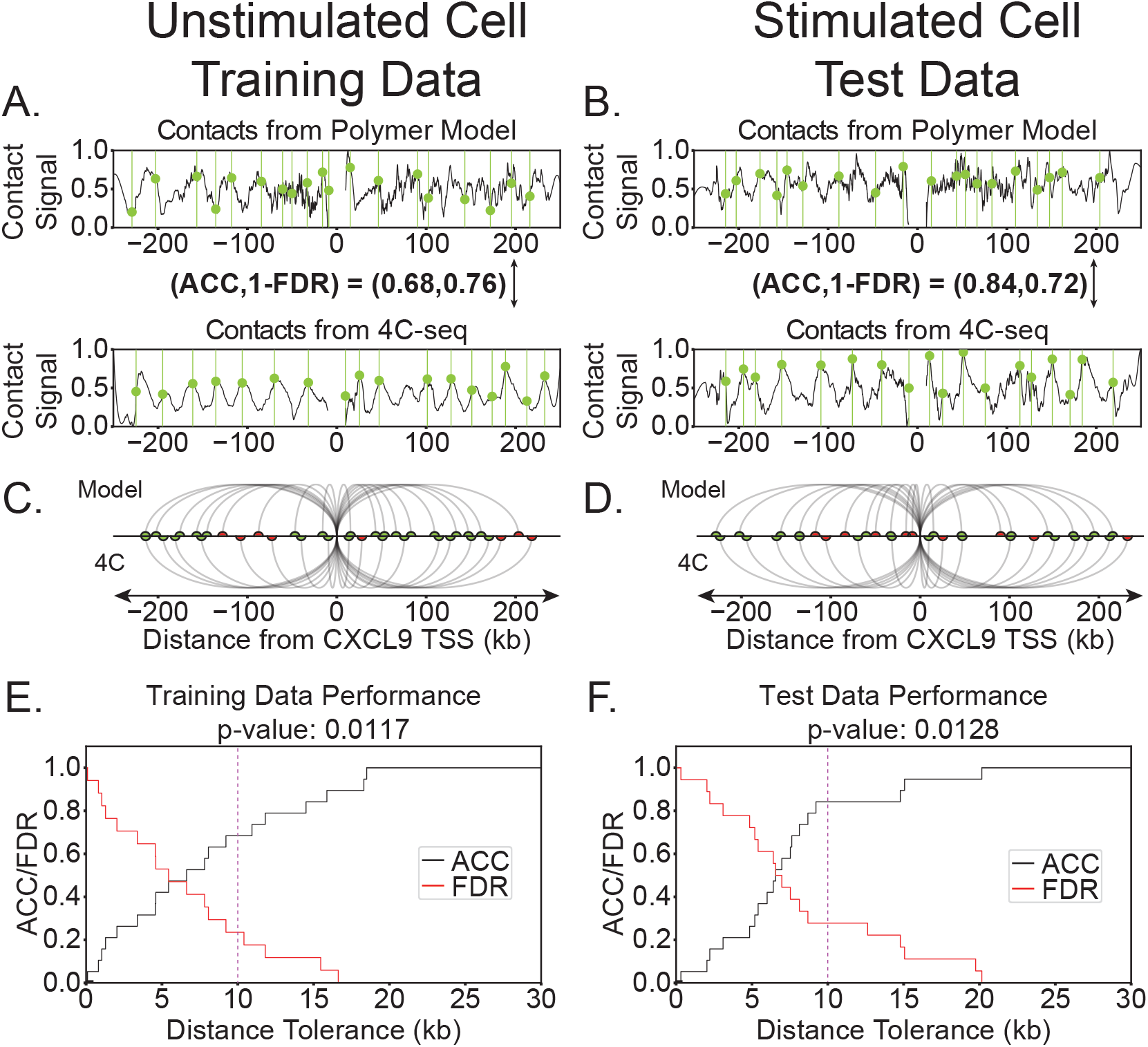
ATAC-EM-informed polymer model predicts close contacts observed in 4C-seq. **A-B**. Contact signals for unstimulated, pre-LPS human macrophage (left) and post-LPS 1hr stimulated macrophage (left). Tracks from the ATAC-EM-informed model (top) and 4C (bottom) are shown with peaks overlayed in green. **C-D**. Predicted contact agreement between our model and 4C. Arcs represent identified contacts. Semicircles represent a 10kb contact, and are colored green if both 4C and the model are in agreement, and are colored red otherwise. **E**. Training set performance. The pre-LPS dataset was used only to train the peak-finding algorithm (all biophysical parameters were fixed). Vertical dashed line shows where accuracy and precision were measured for hyperparameter optimization. (The *p*-value 0.0117 is shown but should not be interpreted as a hypothesis test, since it is on the training set.) **F**. Validation set performance. Within a threshold of 10kbp, the model predicts peaks with an accuracy of 0.84 and a precision (1-FDR) of 0.72 (vertical dashed line). This corresponds to a *p*-value 0.0128 compared to randomly guessing the locations of the same number of peaks.

A straightforward way to assess the predictive power of this model is via two quantities: the proportion of polymer-mechanics-predicted contacts that are within a tolerance distance of a 4C contact (that is, the accuracy of the model); and the proportion of 4C contacts that are within a tolerance distance of a polymer-mechanics-predicted contact (that is, the precision of the model, also called positive predictive value, and is 1 *−* FDR where FDR is the false discovery rate). These are shown in Figure 5EF for various tolerance distances. At high tolerance, prediction becomes perfect, as expected. At the genomic distance of 10kbp, roughly the size of larger gene regulatory elements (37; 38; 39), the accuracy is 0.84 and precision is 0.72. This value has a *p*-value of 0.0128 compared to randomly guessing the locations of the same number of peaks. While this *p*-value is striking, it is comparable to the predictive power of nucleosome positioning for genomic domains on 40 kbp regions in budding yeast (14).

This striking result indicates that polymer mechanics and nucleosome placement, alone, can account for a statistically significant proportion of contacts between two loci. Furthermore, we performed a similar analysis using one 4C replicate to predict another 4C replicate, and found that the average predictive power of 4C is 0.63 (Figure 6B and Supplemental Figure S14 and S15), which is below the predictive power of our model. This between-replicate variability in 4C-seq data is roughly consistent with previous studies and 4C-seq peak-finding algorithms (40; 41; 30). For example, a crude estimate from Figure S1b,c in Geeven et al. (41), their peak identification has a between-replicate accuracy (1-[false positive rate]) of around 70%, roughly in-line with ours. This suggests that the predictive test of our model is not limited by the model itself, but the variability of the 4C data.

**Fig. 6:**
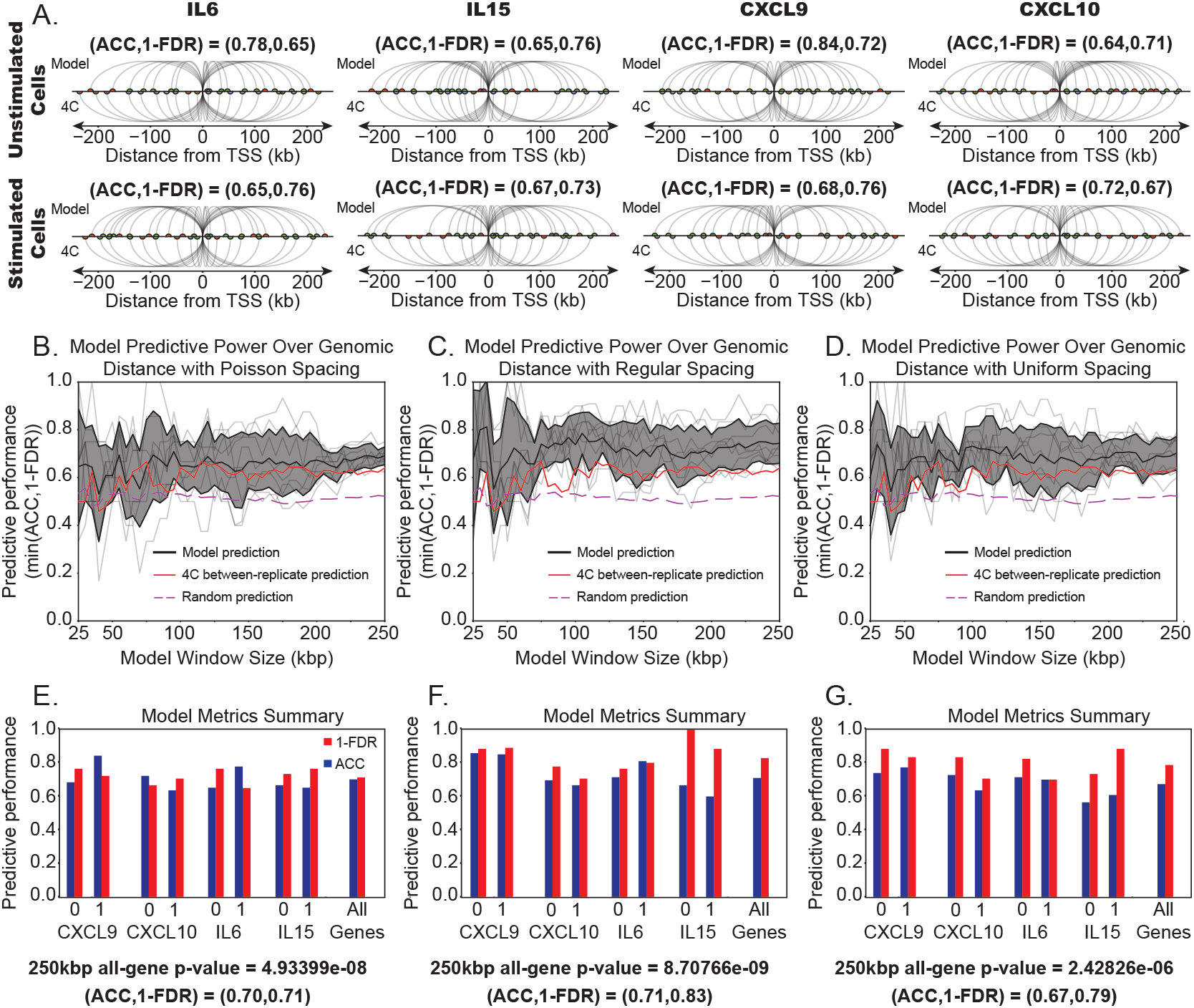
ATAC-seq is predictive of genomic contact across multiple genes and genomic length scales. **A**. Example of predicted contacts for 4C and our polymer model for IL6, IL15, CXCL9, and CXCL10. All contacts were predicted using CXCL9 hyperparameters as described in Figure 5. Contact signals, such as those in 5AB are plotted for all four genes in Figure S16. **B-D**. Performance over genomic distance for all genes for exponentially (B), regularly (C), and uniformly (D) distributed spacing of nucleosomes in low coverage regions. Gray band about model prediction depicts one standard deviation from the average prediction metric. Average 4C metric and random guessing metric are shown for reference. Model overall shows little trend over genomic distance, we find a slight downtrend for small window sizes. At large genomic distance, model performs significantly better than the random predictor with a p-value 4.9399 *×* 10^*−*8^. Performance of the model is similar regardless of model choice for the low-coverage regions. All predicted contacts and contact signals are given in Figures S16-S19. **E-G**. Summary of accuracy and precision metrics for each gene, for both unstimulated (0 hour) and LPS-stimulated (1 hour) human macrophages, and the combination of all four genes being predicted simultaneously for exponentially (E), regularly (F), and uniformly (G) distributed spacing of nucleosomes in low coverage regions.

Note that we report a *p*-value in Figure. 5C, but this should not be interpreted as a hypothesis test since it is part of the training set. Note also that we use a false discovery rate instead of a false positive rate, since the latter is less meaningful for identifying ∼ 10 contacts on a region of ∼ 100kbp.

Beyond CXCL9, we next sought to study other inflammatory genes, IL6, IL15 and CXCL10. Crucially, here we do not retrain the peak-finding hyperparameters (and recall there are no fit parameters in the biophysical model), reusing those we found for CXCL9. The resulting tracks are shown in Figure 6A for the exponential model for nucleosome spacing in low-coverage regions. Tracks for the regular spacing and uniform spacing models are shown in Figures S19. We find that for all genes together, our model yields a metric of 0.70, corresponding to a *p*-value 4.9399 *×* 10^*−*8^, and comparable with that average predictive power of 4C as seen in Figure 6B (red) and Supplemental Figures S14 and S15.

### The range of validity of nucleosome-polymer-mechanics modeling

As mentioned, many other factors influence genome organization besides polymer mechanics and kinks due to nucleosome placement. The scaling behavior of contacts exhibits different regimes at different genomic length scales (6; 42). From these previous studies, we speculate that the simple model developed here would lose validity around 400kbp, where other effects such as fractal globule effects, and topological domains organized by regulatory complexes such as cohesins, may dominate. Therefore, we repeated the analysis, including polymer simulation, for a range of genomic domain sizes, and show the metric in Figure 6B, for all 4 genes and 2 stimulation conditions. We compare this to the ability of one 4C replicate to predict another 4C replicate (red curve). As mentioned above, at this level of predictive performance, we expect the polymer model to be limited by the reproducibility of the 4C data. We also compare to the average predictive power of randomly placed contacts (dashed line; at this predictive performance, the *p*-value would be 0.5). We find that the model performs significantly better than random guessing, and comparable to the limitation of the 4C data. Furthermore, this appears to hold throughout a range of genomic distances, to 250kbp, suggesting that the nucleosomes continue to have a significant impact up to 250kbp. Computational limitations prevented us from simulating larger genomic distances.

### Role of linker length distribution in dense regions

The above results raise the question of which aspect of nucleosome placement, specifically, is encoding the close contacts. We repeated the contact prediction pipeline with the two other (counterfactual) linker length distributions we studied in Figure 2, using the mean spacing derived from EM as in Figure 3G. Predicted and 4C-seq contacts are shown in Supplemental Figures S17, S18 and S19. Intriguingly, we find comparable predictive performance, with similar *p*-values (Figure 6 C, D,F, G). This suggests that the long-range contacts are encoded in the sparse regions, plus the mean density in the dense regions, but not in the details of the distribution in the dense regions.

### Polymer-mechanics-predicted contacts harbor actively engaged regulatory factor DNA-binding motifs

Given that long-range genomic contacts are widely thought to contribute to active gene regulation through the co-localization of regulatory factor binding hubs with distantly located gene promoters, whereby this co-recruitment can, e.g., boost or “enhance” expression, we next examined the presence of regulatory binding factor motifs located within said contacts. To do so, we first selected genomic regions that overlapped as both experimentally observed (based on 4C-seq conserved across two biological replicates) and polymer-mechanics-predicted contacts across unstimulated (0hr) and LPS-stimulated (1hr) conditions. Using this strategy, we identified genomic contact regions that were either dynamically regulated (gained or lost) or static during macrophage stimulation (Supplemental Figure S20A). We next used the MotifMap database to quantify the abundances of transcription factor (TF) binding motifs present within our genomic contact regions and found that these regions harbored a total of 243 motifs spanning 40 distinct transcription factors (Supplemental Figure S20B), with the top 3 most abundant being LEF1, YY1, and IRF8. Notably, LEF1 (lymphoid enhancer binding factor 1) was the most abundant motif within our contacts and is known to regulate gene expression through DNA looping mechanisms (43) and its role in 3d genome organization has particularly been implicated in immune regulation of myeloid lineages (44). We also observed overlap with the binding motif of CCCTC-binding factor CTCF, an architectural protein known to mediate intrachromosomal interactions (45) within a contact that was lost upon inflammatory stimulation. We compared the genomic positions of the motifs contained within our contact regions with publicly available data from genome-wide TF-binding assays (ChIP-seq) for TFs LEF1 and CTCF in Figure 7. Indeed we find that several LEF1 motifs and the CTCF motif captured within our contact regions are occupied by actively bound transcription factor proteins directly in macrophages for CTCF (GSM2544247) or cells of closely related lineage origin to our THP-1 cell line (specifically K562 lymphoblast cells) for LEF1 (GSM2828684).

**Fig. 7:**
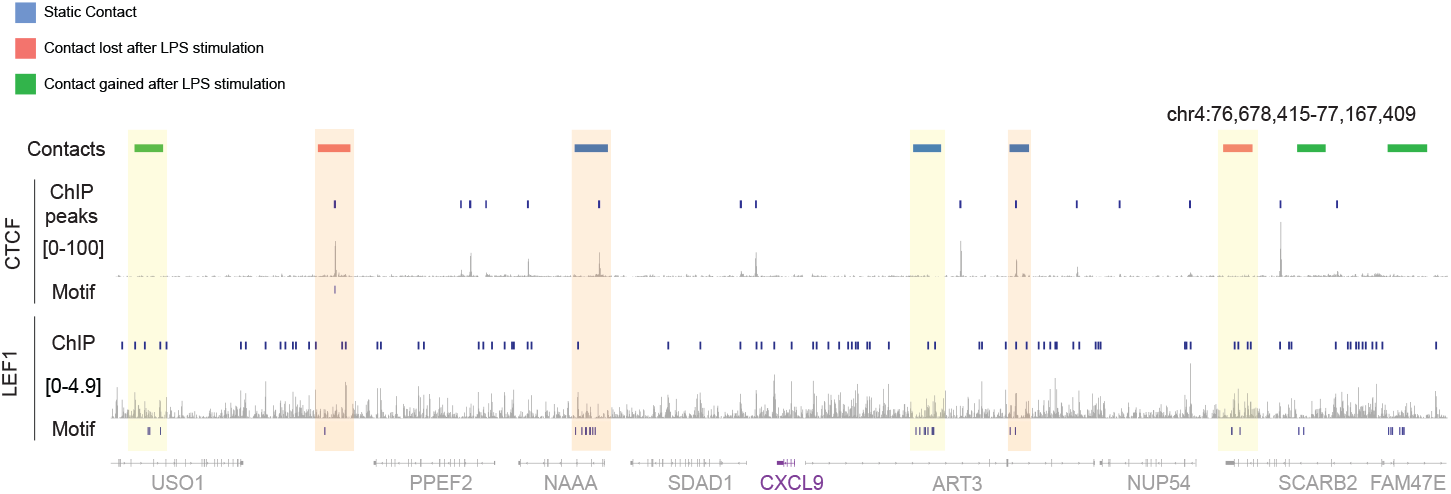
Polymer-mechanics-predicted contacts harbor actively engaged regulatory factor DNA-binding motifs. Genome browser track showing the CXCL9 locus indicating genomic contacts identified in both polymer model and 4C-seq, along with overlapping TF-binding motif for CTCF and LEF1 and the respective ChIP-seq tracks for CTCF binding within macrophages and LEF1 binding within blood lineage-derived K562 lymphoblast cells. Shaded regions indicate contact regions where either a CTCF and LEF1 ChIP-seq peak is present (yellow) or they both coexist (orange).

## Discussion

Besides their role at individual genomic loci, nucleosomes between loci can significantly modulate their long-range contacts. Changing nucleosome density within the physiologically-observed range can lead to order-of-magnitude changes in contact probability. More surprisingly, we find that the specific placement of these nucleosomes between loci, even at approximately the same total number, can introduce or reduce contacts.

The specific contacts predicted by the polymer model, after training on ATAC-seq and electron microscopy data, were statistically significant and on-par with the difference between replicates of 4C-seq. Indeed, 71% of close contacts within four 500 kbp genomic regions, before and after inflammation, were correctly predicted by the model. This is surprising, since many other factors are known to influence 3d genome organization, but is consistent with prior work by Wiese et al. (14) that demonstrate that nucleosome positioning is predictive of other aspects of 3d genome structure in budding yeast at ∼ 40 kbp distances. Wiese et al. (14) speculated that the situation would be different in higher eukaryotes, but here we demonstrate that the role of nucleosomes in encoding 3d genome organization is conserved in the context of human-derived macrophages, and to ∼ 10x longer genomic distances. We therefore add to a growing body of work that suggests nucleosomes dominate, for example, the effects of the presence or absence of cohesins (8), and that cohesins are not always necessary to predict structure (12). To be sure, there is clear evidence that factors besides nucleosomes and polymer mechanics help determine and modulate 3d genome structure. These other factors have been reviewed and categorized as (11): motor and extrusion factors; compartmentalization and phase separation; landmark association (e.g., the nuclear lamina); and topological constraints preventing obedience to ideal polymer physics (e.g., fractal globule effects). And yet, our results suggest that, while genome contacts on 100 kbp scales are multifactorial, some of these contacts may be amenable to mechanistic, physical explanation. A possible partial explanation for the success of the model is that it operates on the short length scale where ideal polymer scaling has been previously observed (11), which also corresponds to the size of typical TADs. Computational cost prevented us from simulating larger genomic sizes. While the algorithm scales linearly with the number of nucleosomes to generate individual polymer conformations, the number of conformations needed to sample increasingly-rare contacts grows as a power-law with genomic distance.

The image analysis and pathfinding algorithm developed herein was necessary, since ATAC-seq coverage is low in most of the genome. Our method, which uses a novel pathfinding algorithm that weighs paths by both their intensity under EM and their path-length, yielded a distribution of nucleosome spacings consistent with previous work (33; 34; 8). It would be intriguing to apply this method to EM data on specific regions of the nucleus, and to cells in different regulatory states.

This model assumed that causality runs from nucleosome placement, to polymer mechanics, to conformation and close contacts. While the predictive success of this model demonstrates this is sufficient to explain a significant fraction of contacts, other mechanisms are possible. For example, it is possible that 3d conformation influences nucleosome placement, since histones may bind more easily to certain DNA conformations. Moreover, the purely causal, mechanistic assumptions of the model may also be a detriment if the goal were singularly to predict 4C data from ATAC data — something that could be better achieved by incorporating more information or removing physical constraints implied by the model.

We emphasize that the granularity of this polymer mechanics model is optimal for this 100-kilo-basepairs-scale inquiry. The configurations are computed using efficient Monte Carlo sampling where analytical expressions (46; 47) are used for the mechanics of unbound DNA, greatly increasing the efficiency of the code. Models with more atomistic detail would be computationally prohibitive, and models on larger genomics scales would miss the effect of individual nucleosomes. This mesoscale approach, and the specific model and pipeline presented here, generalize to other genes, and can be used to address other cases of cells undergoing a state change. Having established the quantitative ability of modeling at this scale, it may be possible to integrate these results into larger-scale models of genome architecture (48; 12), via bias in the angular orientation between coarse-grain simulation beads. In the other direction of scales, molecular factors that directly influence nucleosomes (reviewed in (10)) could be incorporated into this model.

Throughout evolution, the sequence structure of the genome is undoubtably shaped through selective pressures associated with physical polymeric properties of the genome (49). However the intricacies of this co-evolution of sequence and physical property remain largely undefined. Notably, there are long-standing observations that TF binding sites occur in relatively high-density clusters (or hotspots) throughout the genome (50; 51). Furthermore, the positioning of nucleosomes is heavily guided by genetic sequence yet still subject to precise nucleosome eviction events dynamically regulated according to cell state (52). Our findings suggests a new paradigm through which nucleosome placement might favor contact-(or loop-) formation between regulatory binding sites with gene promoters, thereby influencing transcription factor search dynamics (53).

## Data Availability

The data underlying this article are available at GEO (www.ncbi.nlm.nih.gov/geo) under accession number GSE278886. Software to simulate the polymer model is available at github.com/ajspakow/wlcstat and github.com/jcorrett/ATAC-EM_Polymer_Model.

## Author contributions statement

J.C. wrote software pipeline, performed all computational simulations and data analysis. H.S., J.L., and P.K.V. performed experiments and data collection. J.C., A.S., T.L.D., and J.A. conceptualized the project and supervised the work. J.C. and J.A. wrote the manuscript.

## Acknowledgments

We thank Elizabeth Read (UCI) for valuable discussion. This work is funded by NSF DMS 2052668, NSF Emerging Frontiers URoL 2022182, NIH T32 GM136624, NSF DMS 1763272, and the Simons Foundation (594598, QN).

## Supplementary Material

**Figure S1:**
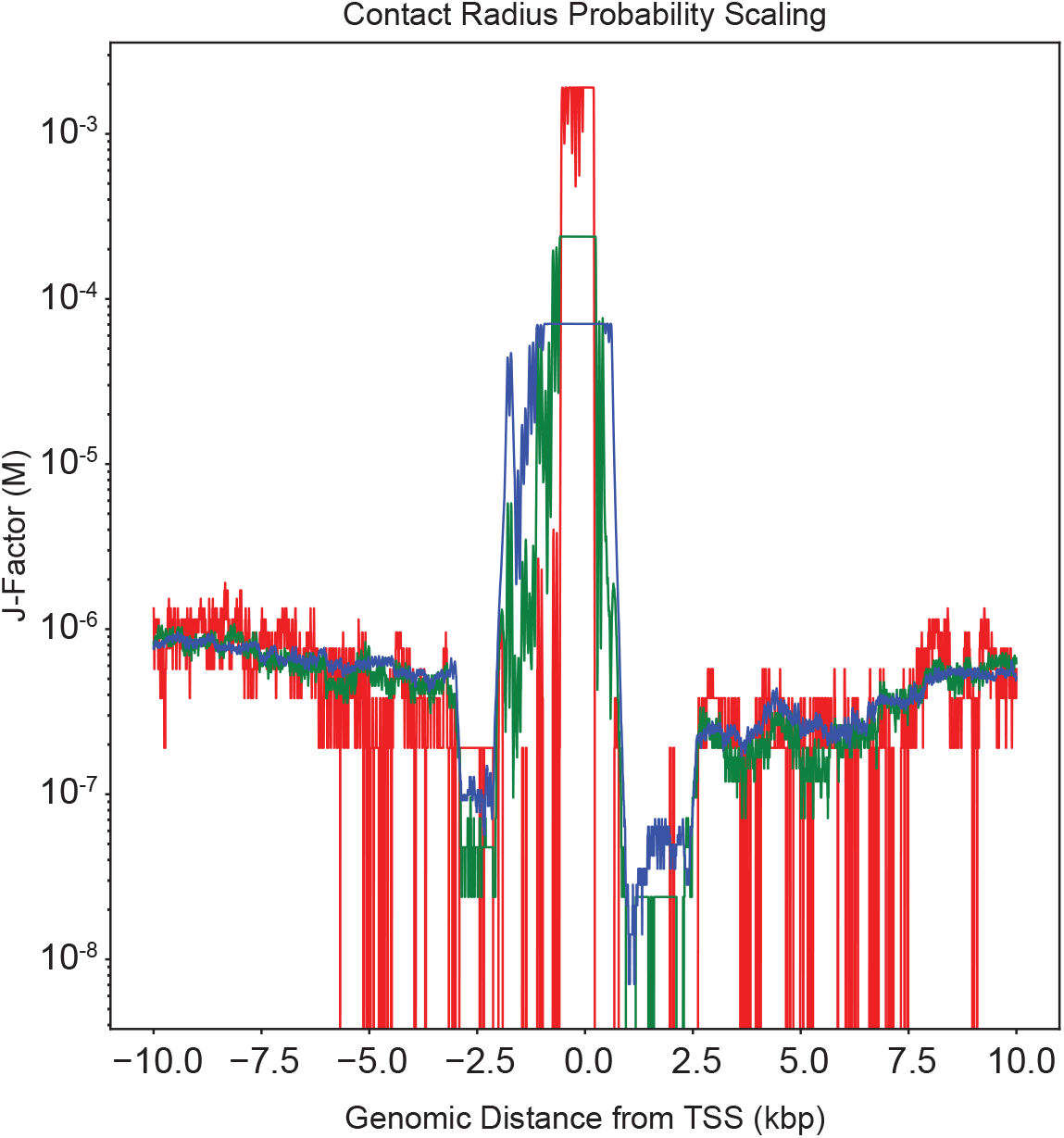
Contact probability scales directly with the contact radius. Shown for a contact radius of 5, 10, and 15 nm (red, green, and blue respectively). Units are in J-factor, which is contact probability scaled by a contact volume of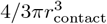.

**Figure S2:**
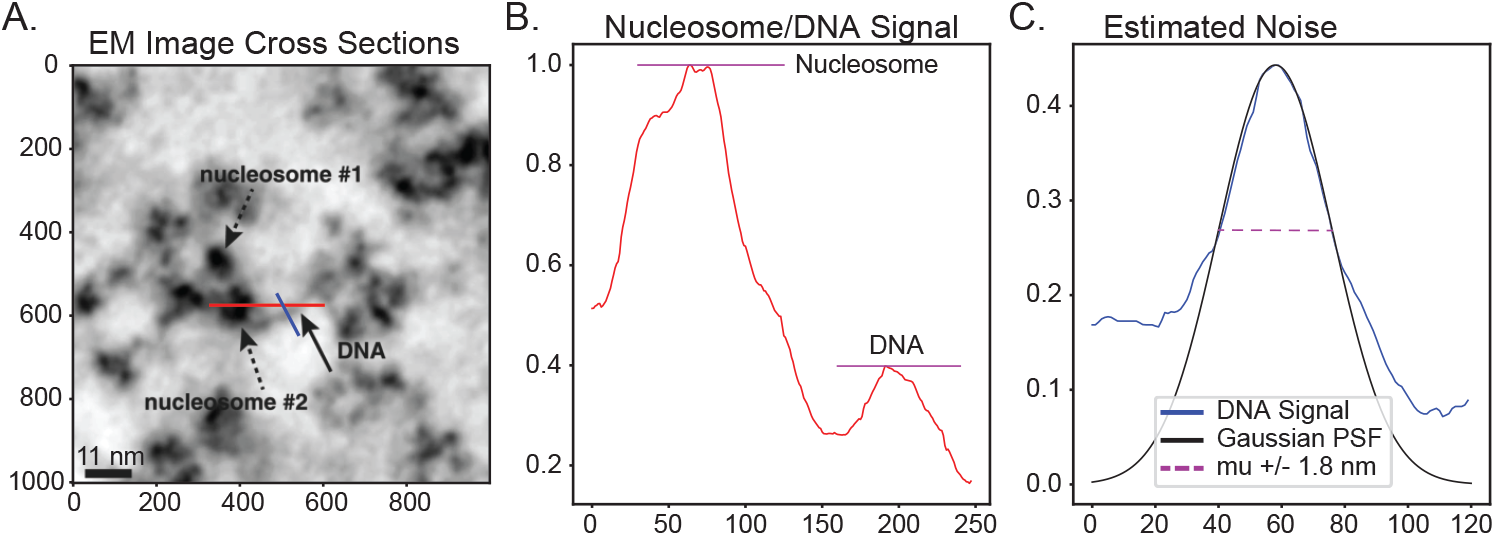
Estimated EM properties from ChromEMT. **A**. ChromEMT sample with identified nucleosomes and DNA (Figure 3C, Ou et al. [19]). **B**. Red image profile line from A. crossing a nucleosome and DNA. Nucleosomes are approximately 2.5 times higher in signal than DNA. **C**. Blue image profile line from A., as a cross section of DNA. Fit Gaussian point spread function (black) indicates a gaussian blur level of 1.8 nm. Mean plus or minus 1.8 nm shown in magenta.

**Figure S3:**
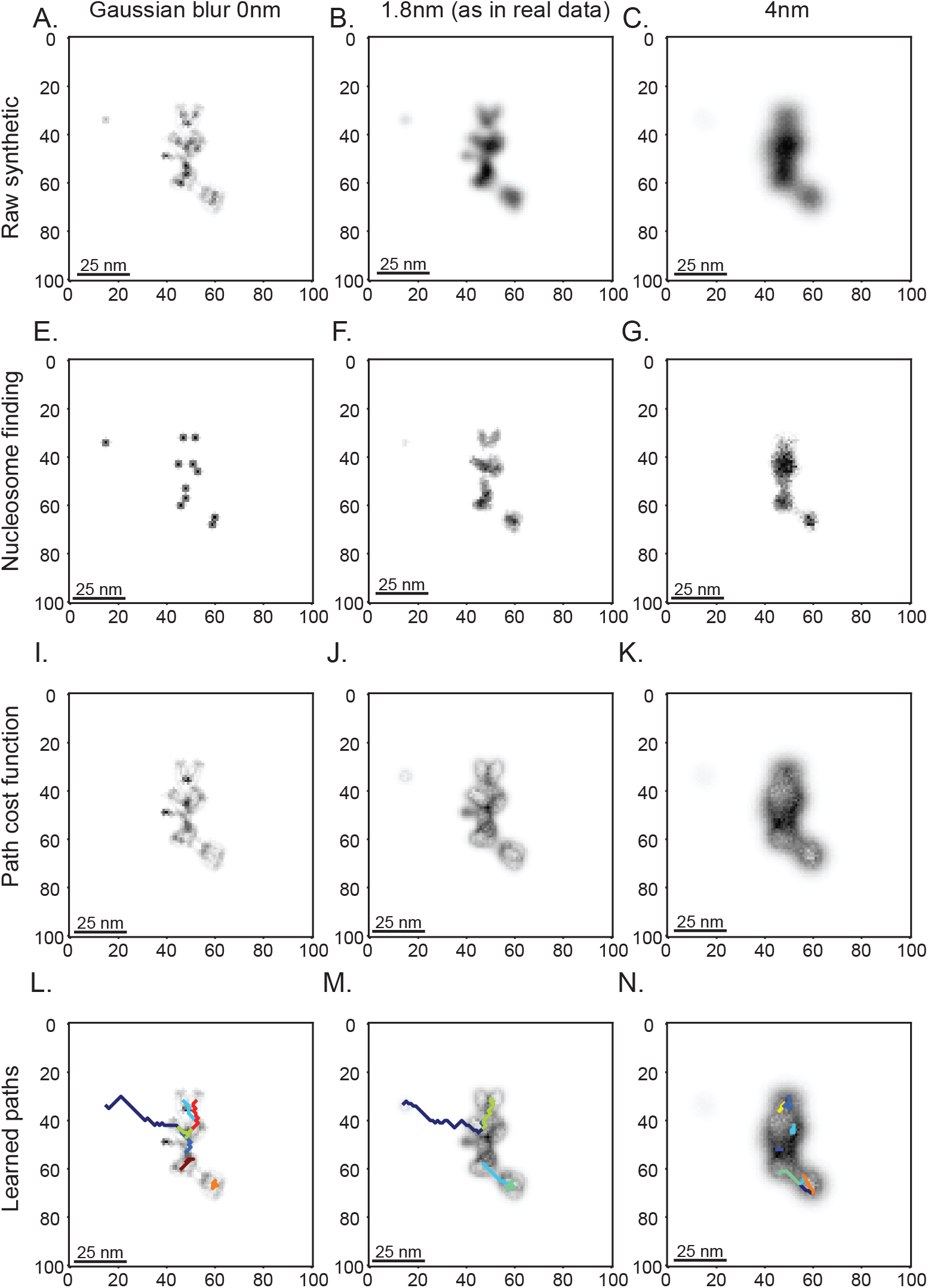
Synthetic EM nucleosome spacing pipeline visualization for mean nucleosome spacing of 30 basepairs. **A-C**. Raw synthetic EM images for Gaussian noise of 0.0 nm (left), 1.8 nm (middle), and 4.0 nm (right). Nucleosomes are 2.5 times darker than DNA as described in Methods and Figure S2B. **E-G**. Identified nucleosomes after image binarization. **I-K**. Images fed into Dijkstra’s algorithm as the cost graph after nucleosomes identified in E-G are subtracted from the raw EM images. **L-N**. Sample DNA linkers identified by Dijkstra’s algorithm.

**Figure S4:**
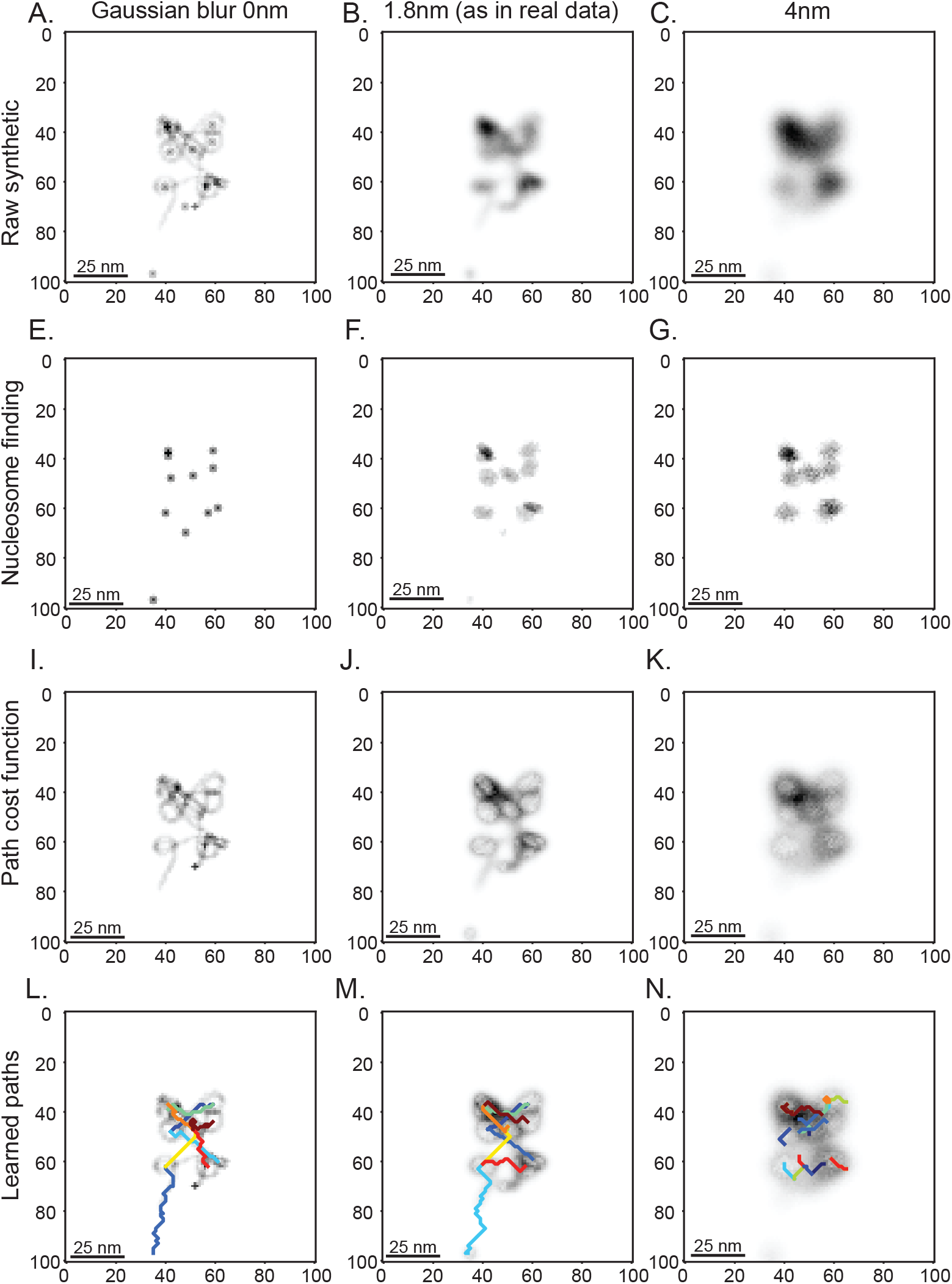
Synthetic EM nucleosome spacing pipeline visualization for mean nucleosome spacing of 60 basepairs. **A-C**. Raw synthetic EM images for Gaussian noise of 0.0 nm (left), 1.8 nm (middle), and 4.0 nm (right). Nucleosomes are 2.5 times darker than DNA as described in Methods and Figure S2B. **E-G**. Identified nucleosomes after image binarization. **I-K**. Images fed into Dijkstra’s algorithm as the cost graph after nucleosomes identified in E-G are subtracted from the raw EM images. **L-N**. Sample DNA linkers identified by Dijkstra’s algorithm.

**Figure S5:**
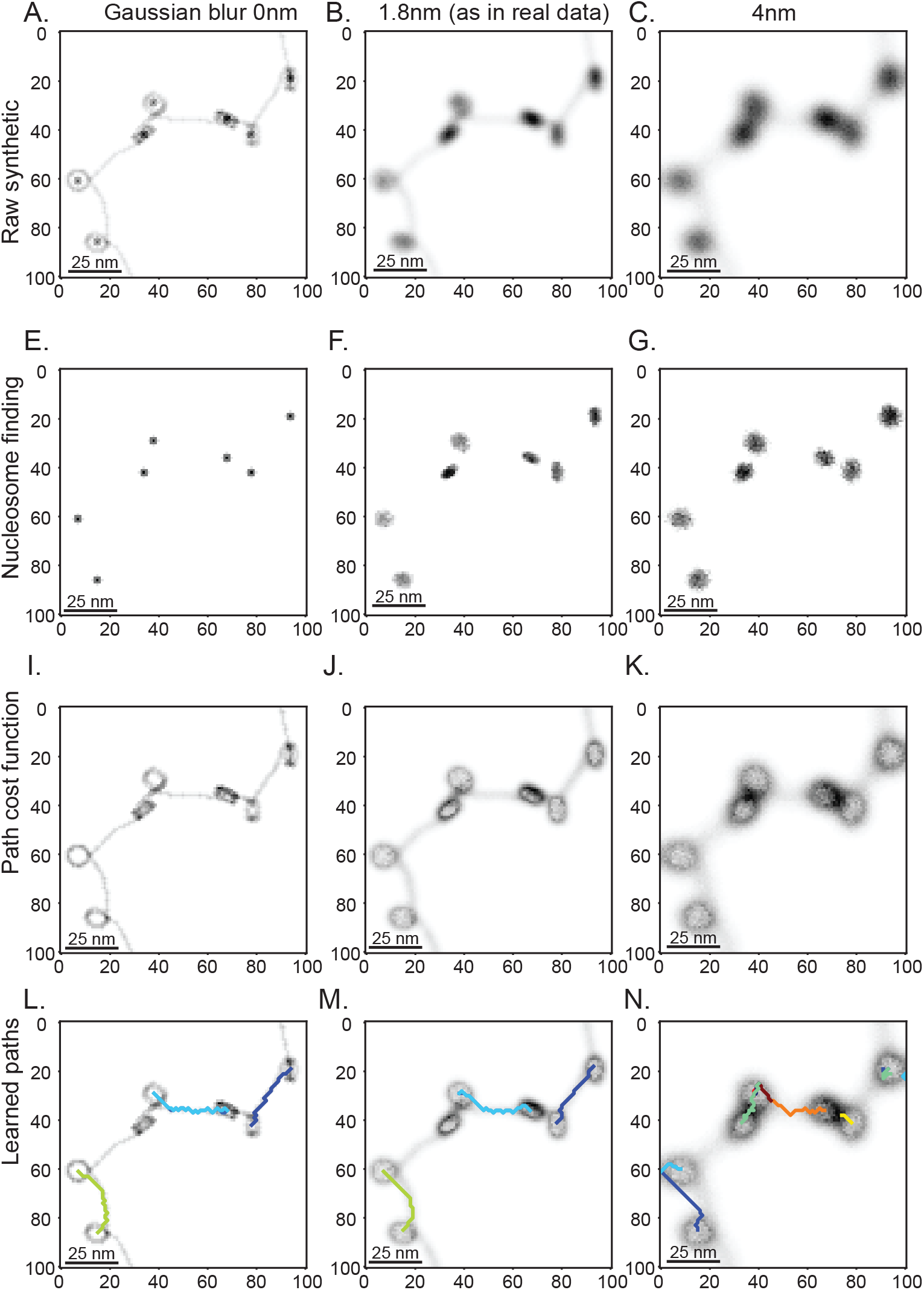
Synthetic EM nucleosome spacing pipeline visualization for mean nucleosome spacing of 90 basepairs. **A-C**. Raw synthetic EM images for Gaussian noise of 0.0 nm (left), 1.8 nm (middle), and 4.0 nm (right). Nucleosomes are 2.5 times darker than DNA as described in Methods and Figure S2B. **E-G**. Identified nucleosomes after image binarization. **I-K**. Images fed into Dijkstra’s algorithm as the cost graph after nucleosomes identified in E-G are subtracted from the raw EM images. **L-N**. Sample DNA linkers identified by Dijkstra’s algorithm.

**Figure S6:**
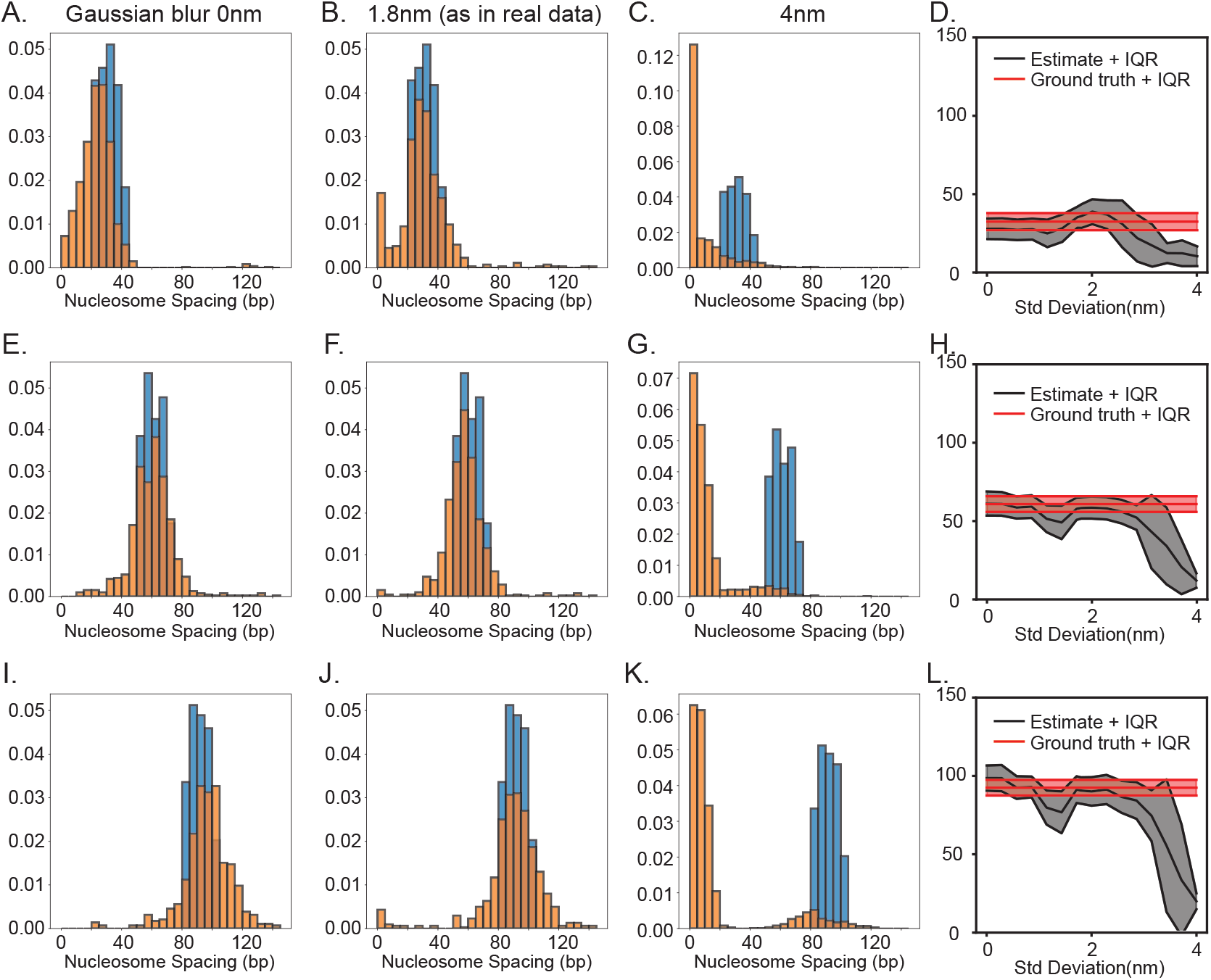
Estimated nucleosome spacing for different mean spacing and Gaussian blurs. **A-C**. Learned nucleosome spacing histograms for a mean of 30 basepairs and a Gaussian blur of 0.0 (left), 1.8 (middle), and 4.0 (right) nm. **D**. Parameter sweep of Gaussian blur intensity for sampling nucleosome spacing from synthetic EM for a mean nucleosome spacing of 30 bp. **E-G**. Learned nucleosome spacing histograms for a mean of 60 basepairs and a Gaussian blur of 0.0 (left), 1.8 (middle), and 4.0 (right) nm. **H**. Parameter sweep of Gaussian blur intensity for sampling nucleosome spacing from synthetic EM for a mean nucleosome spacing of 60 bp. **I-K**. Learned nucleosome spacing histograms for a mean of 90 basepairs and a Gaussian blur of 0.0 (left), 1.8 (middle), and 4.0 (right) nm. **L**. Parameter sweep of Gaussian blur intensity for sampling nucleosome spacing from synthetic EM for a mean nucleosome spacing of 90 bp.

**Figure S7:**
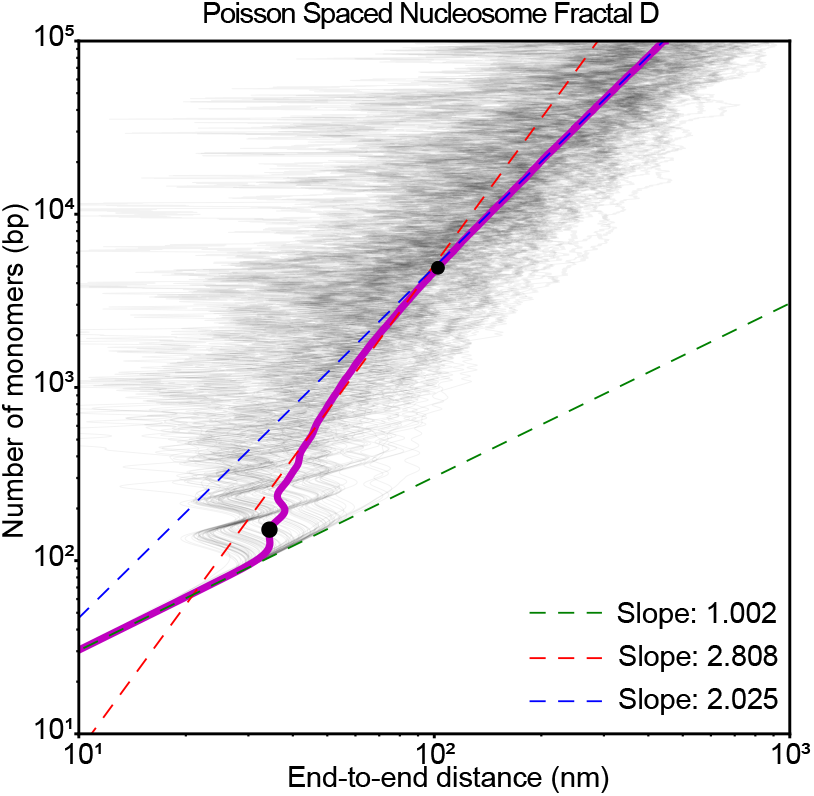
Estimation of the chromatin packing scaling factor *D*. 10,000 polymers of length 10 kbp are simulated with nucleosomes placed randomly according to the exponential distribution derived in Figure 3G. *D* is estimated by a log-log linear fit between the persistence length of DNA (≈ 150*bp*) and the upper bound *r* = 102.4 (black points) as described in Li et al. [35]. To the left of this region, polymers extend linearly with a log-log linear slope ≈ 1, and to the right polymers extend as a random walk with log-log linear slope ≈ 2 in accordance with standard behaviour of Gaussian polymer chains.

**Figure S8:**
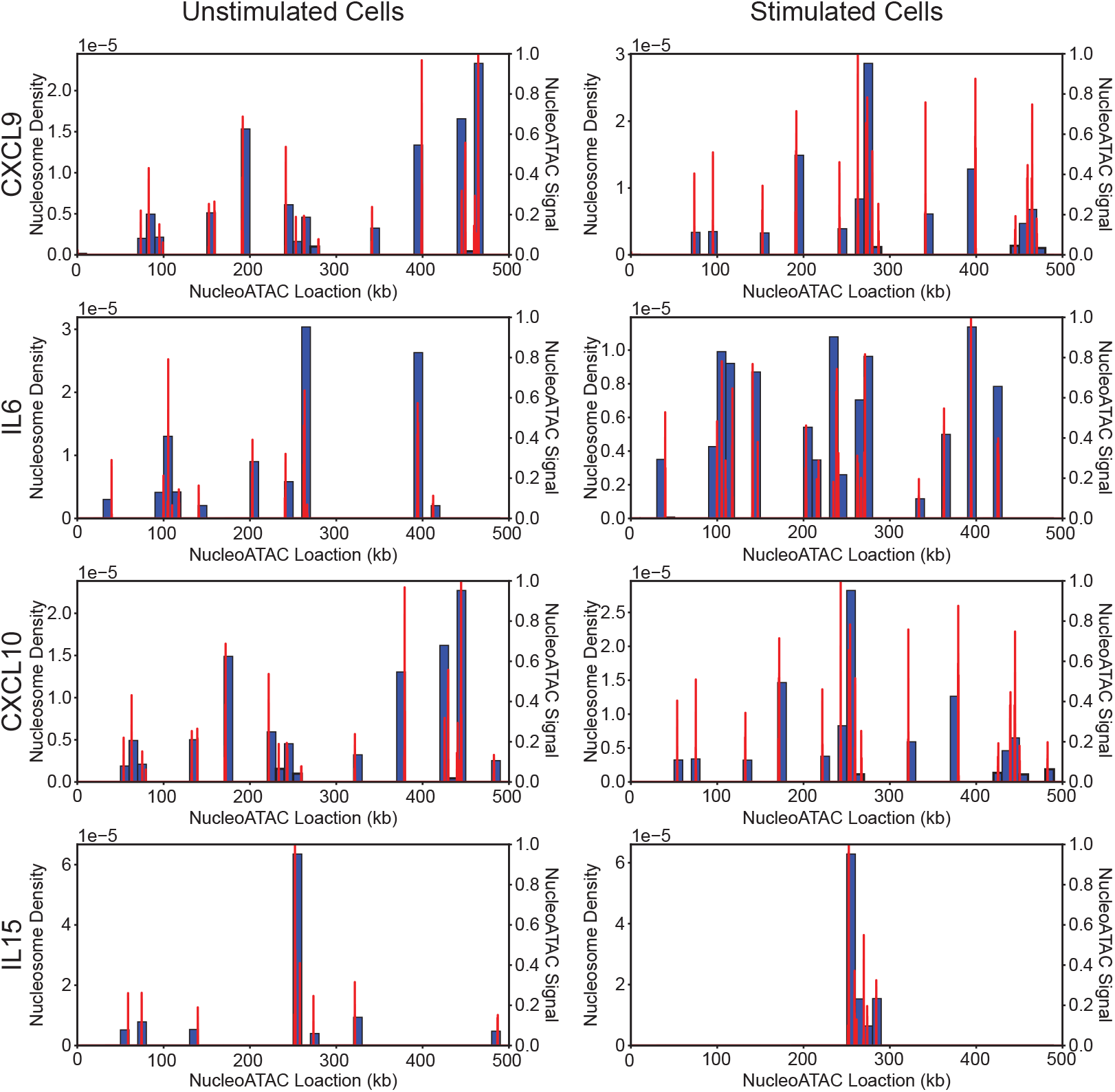
NucleoATAC score normalized to maximum peak height (red, right axis) and nucleosome algorithm placement densities (blue, left axis) for CXCL9, IL6, CXCL10, and IL15 for ATAC-seq data from both unstimulated (left) and stimulated macrophages (right). Peaks in nucleosome density correspond with NucleoATAC peaks and are in general correlated with NucleoATAC peak height.

**Figure S9:**
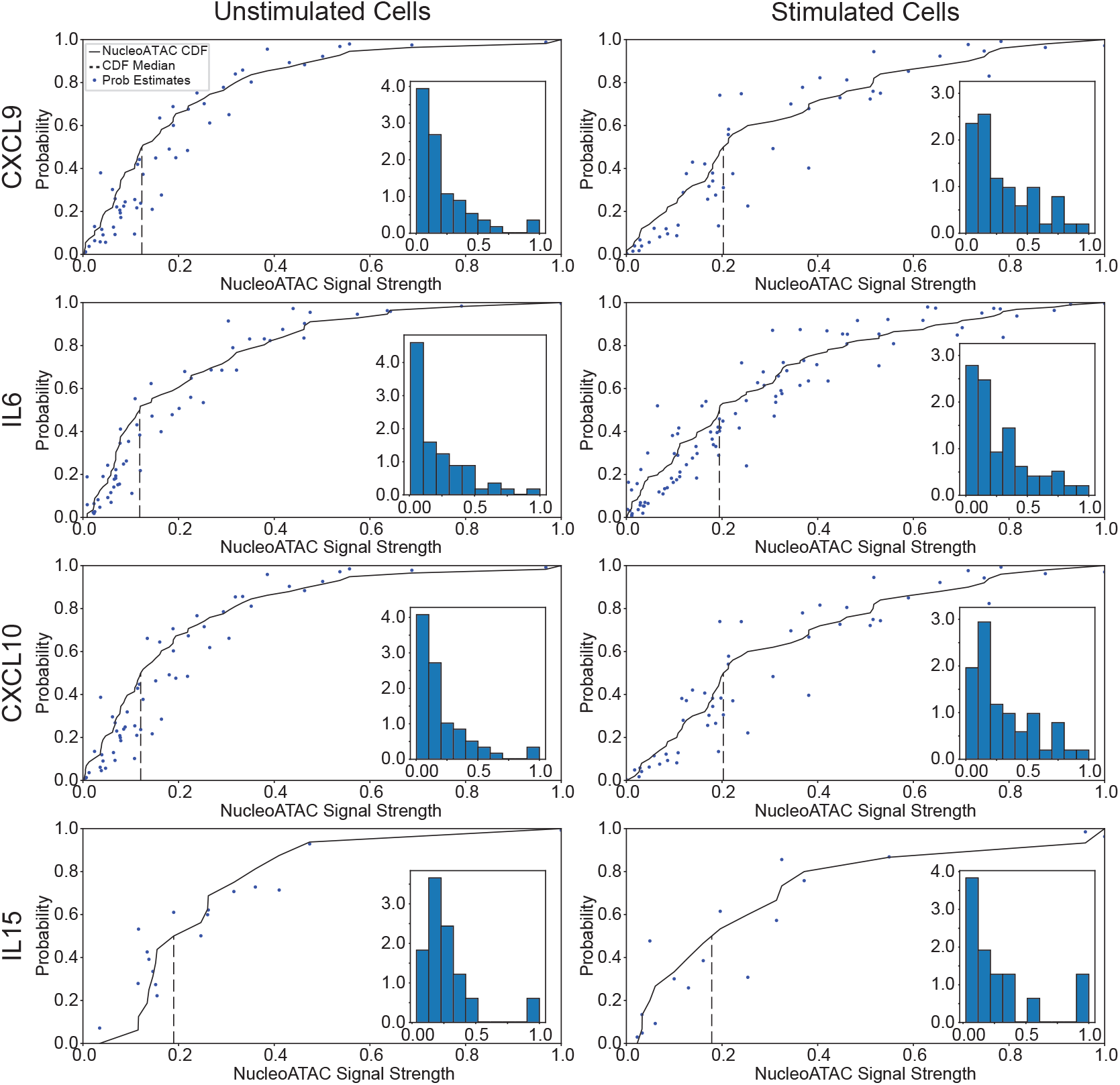
Nucleosome placement probability (blue) and NucleoATAC cumulative distributions (black) as functions of normalized NucleoATAC signal strength for all genes and LPS stimulation time points. Medians of CDF indicated as dashed vertical lines. Inset histograms of NucleoATAC signal value at the peak regions. This data is used to compute CDFs. In general, the probability of nucleosome placement as a function of NucleoATAC signal strength agrees with the distribution of NucleoATAC peaks.

**Figure S10:**
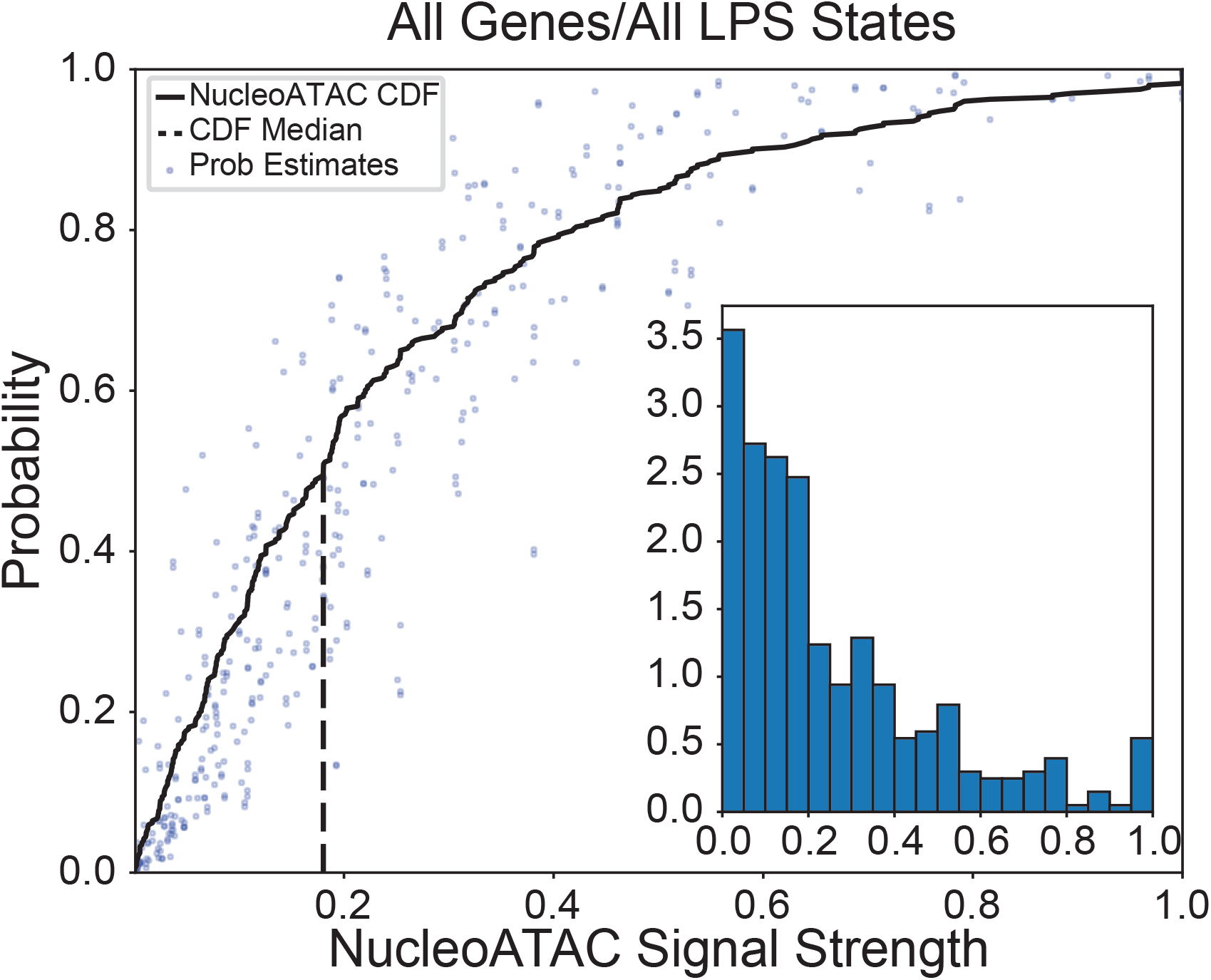
Same as Figure S9, but with all eight data sets combined as an ensemble.

**Figure S11:**
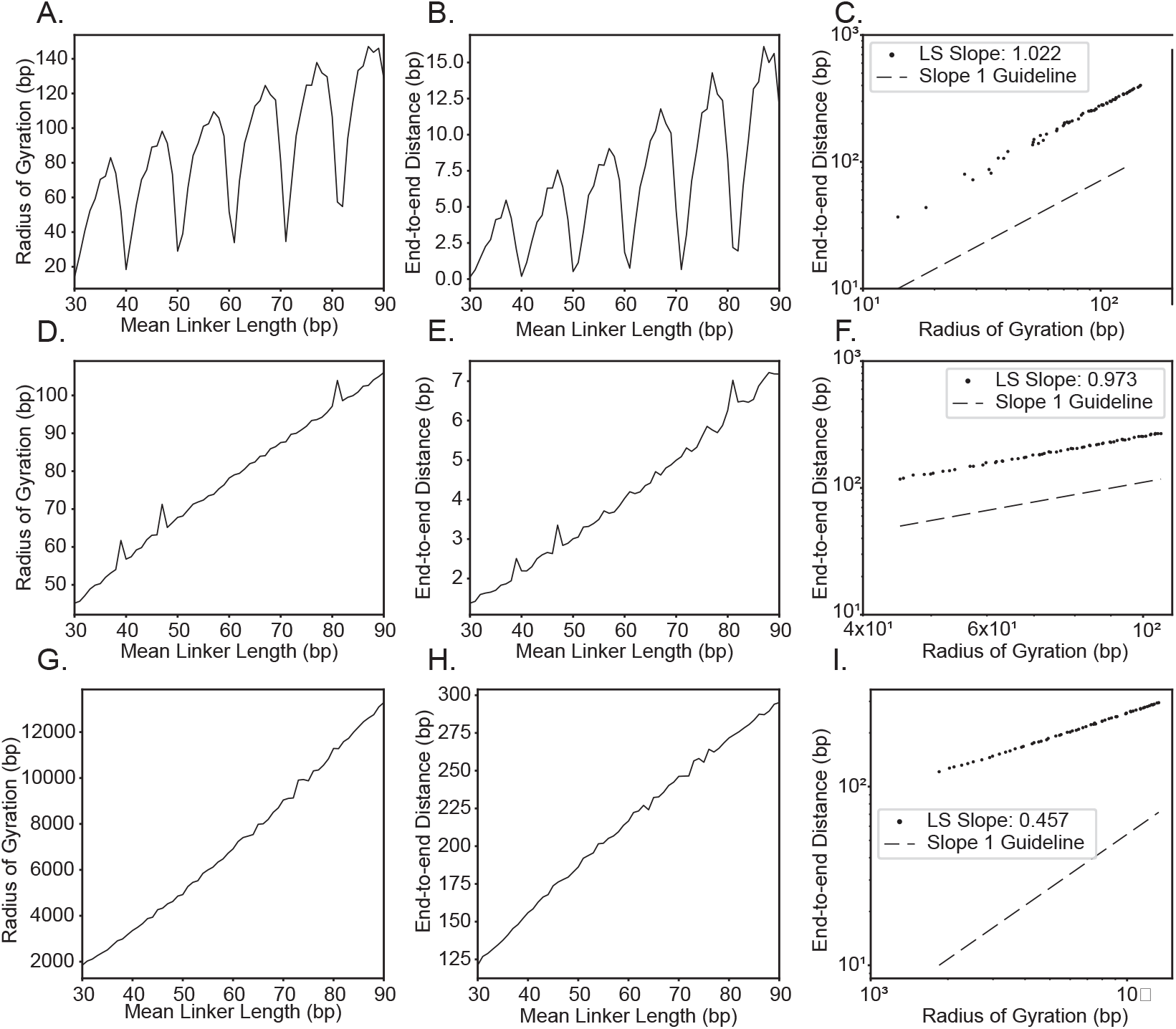
Polymer statistic analysis of CXCL9 simulated using twistable worm-like chain with nucleosomes placed from learning algorithm. Radius of Gyration (RoG) versus End-to-end distance as a function of nucleosome spacing. **A-C**. Mean nucleosome spacing versus radius of gyration for constant nucleosome spacing with *±*0 random spread. **D-F**. Mean nucleosome spacing versus radius of gyration for constant nucleosome spacing with *±*10 random spread. **G-I**. Mean nucleosome spacing versus radius of gyration for exponentially distributed nucleosome spacing.

**Figure S12:**
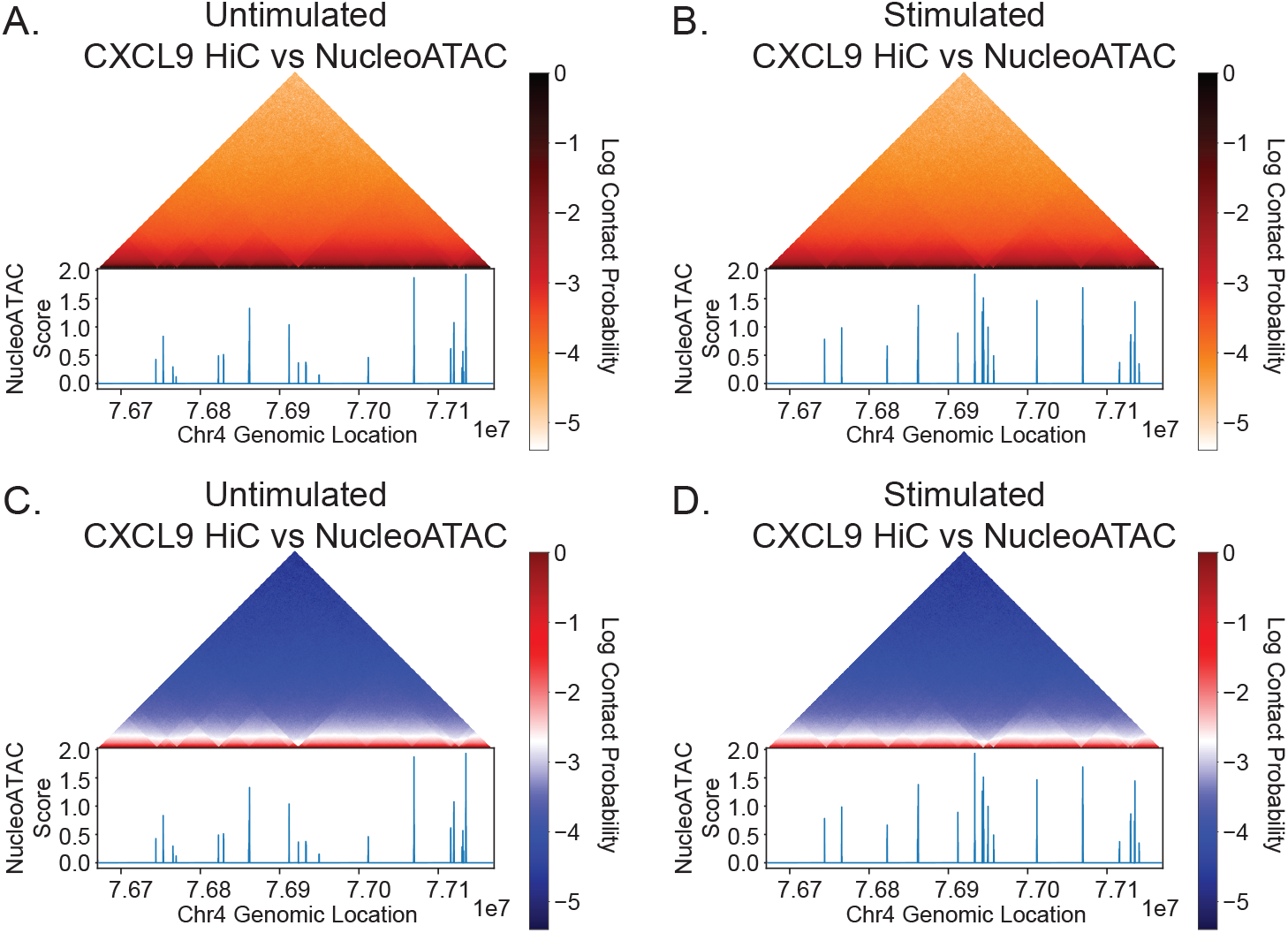
**A**. (Top) Average pairwise contact map for *±*250 kbp of the CXCL9 TSS in the unstimulated model. (Bottom) NucleoATAC signal for *±*250 kbp of the CXCL9 TSS in unstimulated macrophages. Edges of association domains coincide with peaks in the NucleoATAC signal. **B**. Same as A, but after LPS stimulation. Edges of association domain again correspond to (post-stimulation) NucleoATAC peak locations. **C-D**. Same as A and B, but with a different heatmap to better visualize the structure in the data.

**Figure S13:**
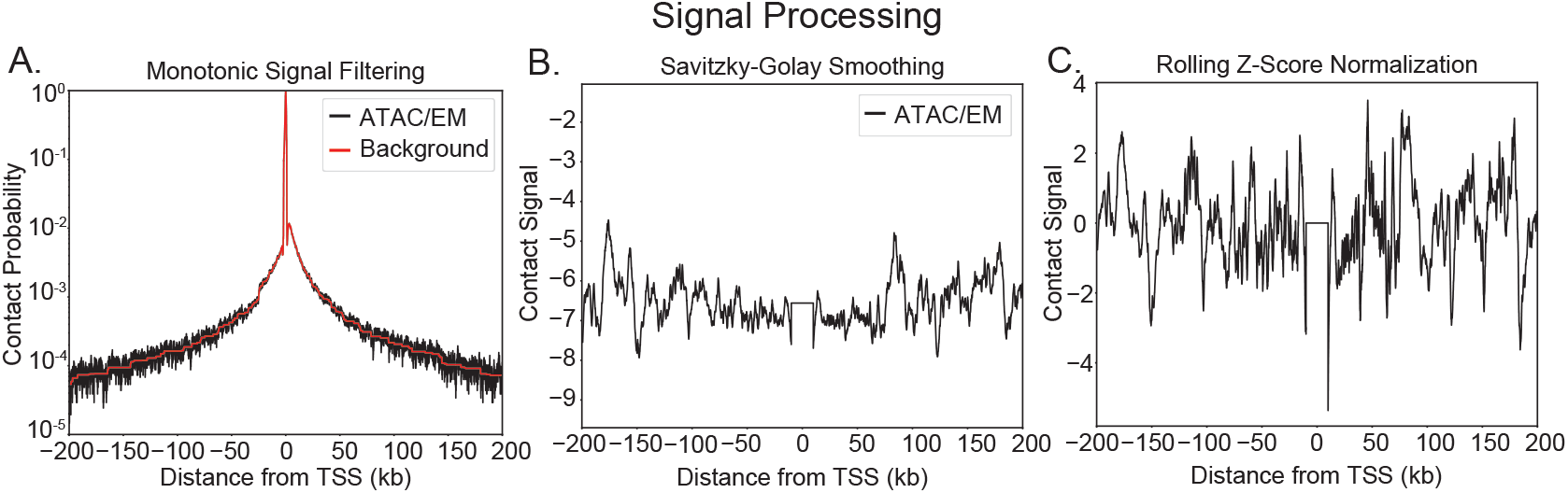
Close contact prediction pipeline. Processed contact signal after smoothing and filtering (see Methods).

**Figure S14:**
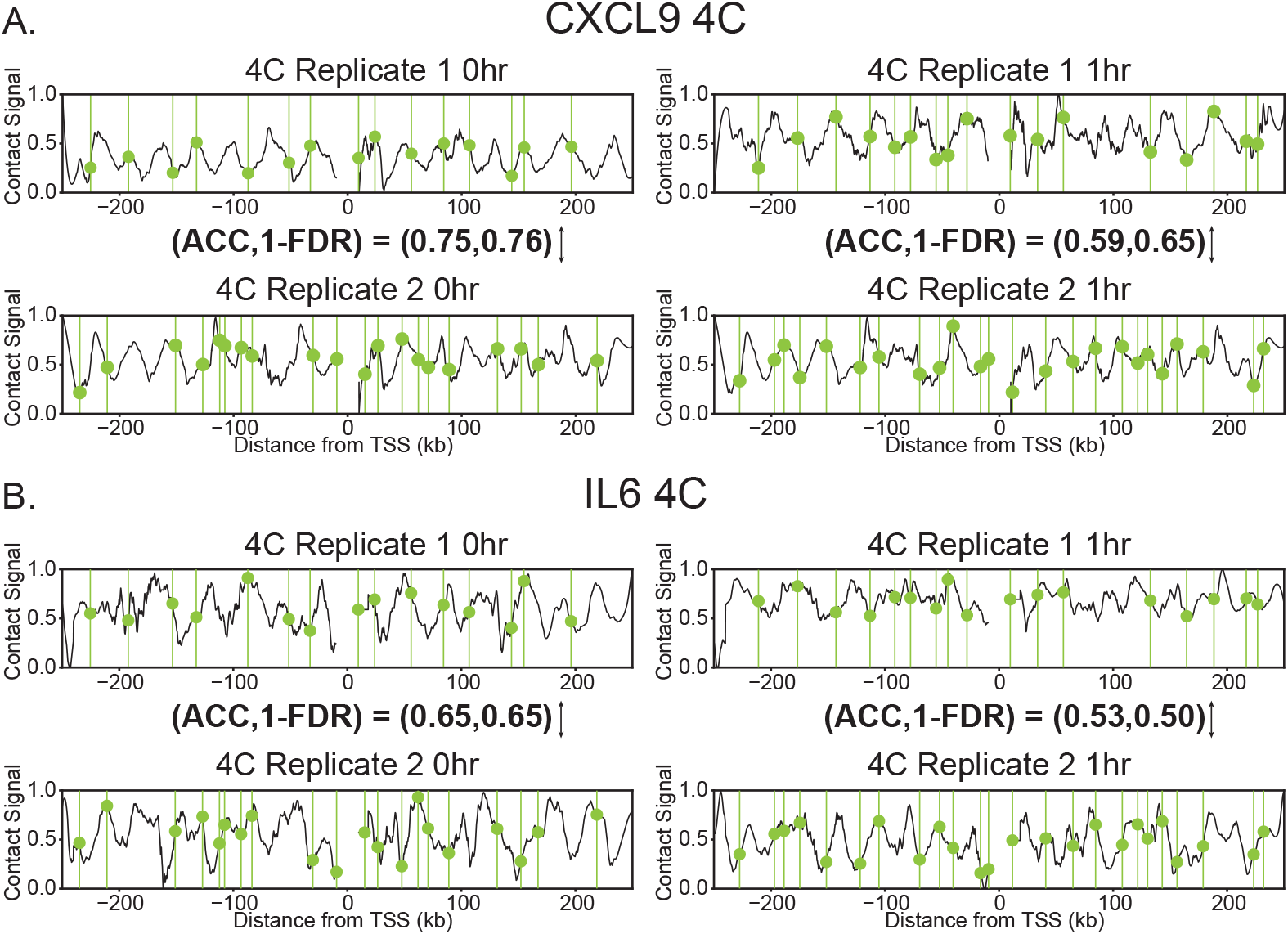
Close contact discovery in two replicates of 4C-seq for CXCL9 and IL6. One replicate is used to predict the close contacts in the other. Hyperparameters of the wavelet peakfinding algorithm are optimized to maximize the minimum of accuracy and precision, same as in the polymer model. By 1 hour post-LPS stimulation, the THP-1 macrophage population is undergoing a dynamic process, where we expect significant heterogeneity. Indeed, we observe that the reproducibility between replicates drops for 1-hour post-LPS stimulation compared to “0hr”, i.e., unstimulated cells.

**Figure S15:**
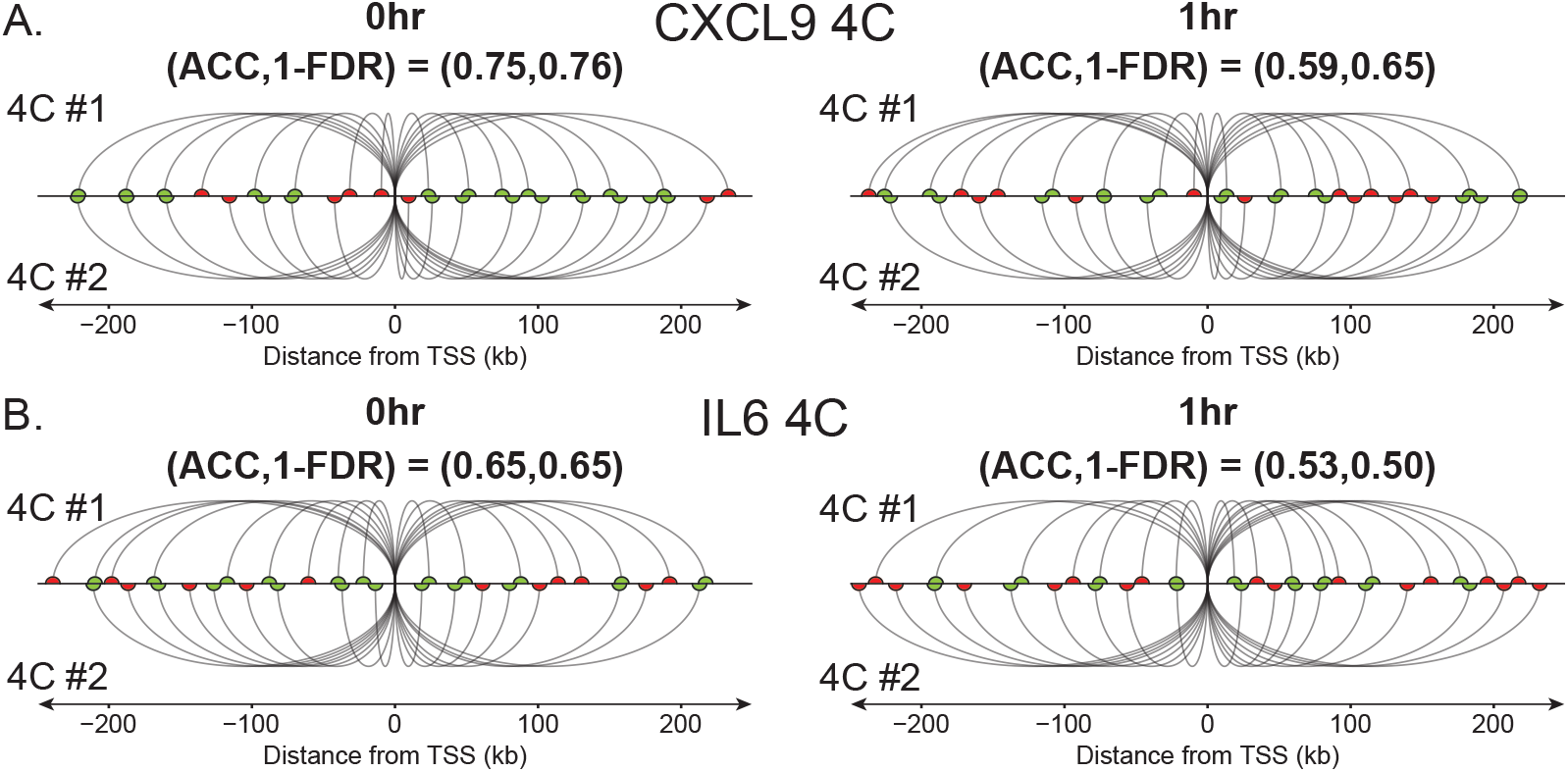
Predicted contacts for data in Supplemental Figure S14

**Figure S16:**
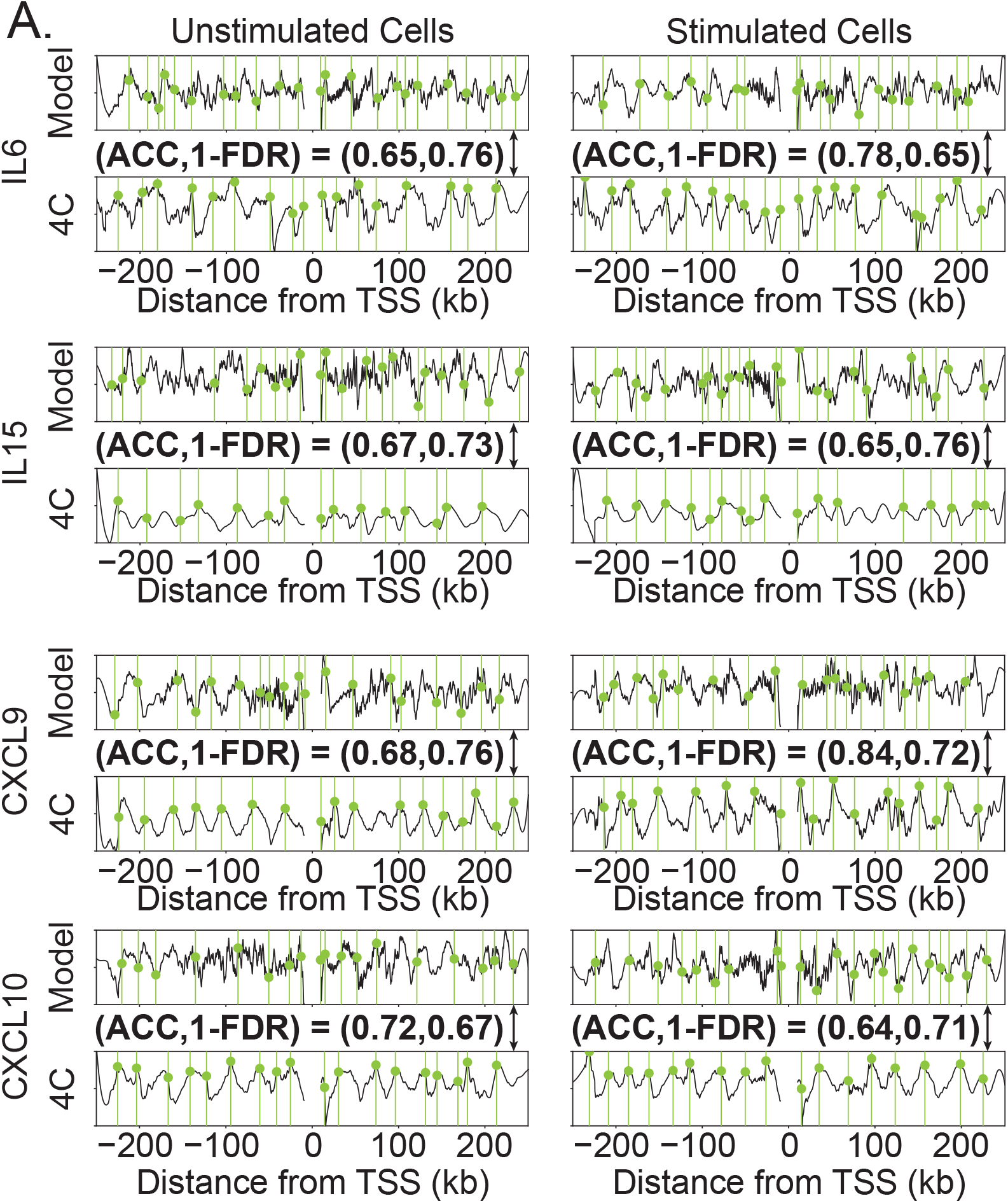
Contact signals for unstimulated, pre-LPS human macrophage (left) and post-LPS 1hr stimulated macrophage (left) for IL6, IL15, CXCL9, and CXCL10. Tracks from the ATAC-EM-informed model (top) and 4C (bottom) are shown with peaks overlayed in green. Spacing of nucleosomes in low coverage regions are distributed exponentially, as described by the model in Figure 2CF.

**Figure S17:**
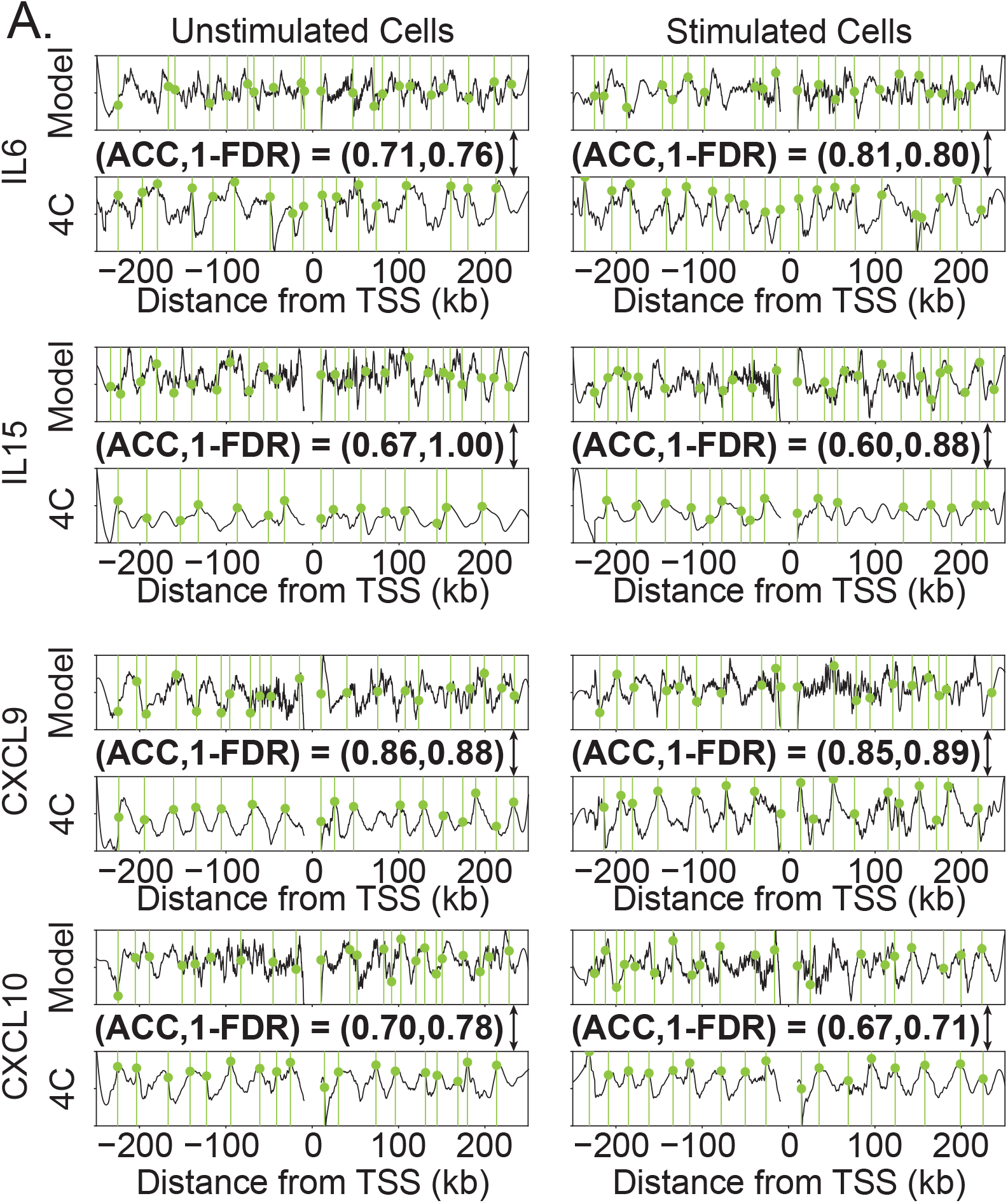
Contact signals for unstimulated, pre-LPS human macrophage (left) and post-LPS 1hr stimulated macrophage (left) for IL6, IL15, CXCL9, and CXCL10. Tracks from the ATAC-EM-informed model (top) and 4C (bottom) are shown with peaks overlayed in green. Spacing of nucleosomes in low coverage regions are distributed evenly, as described by the model in Figure 2AD.

**Figure S18:**
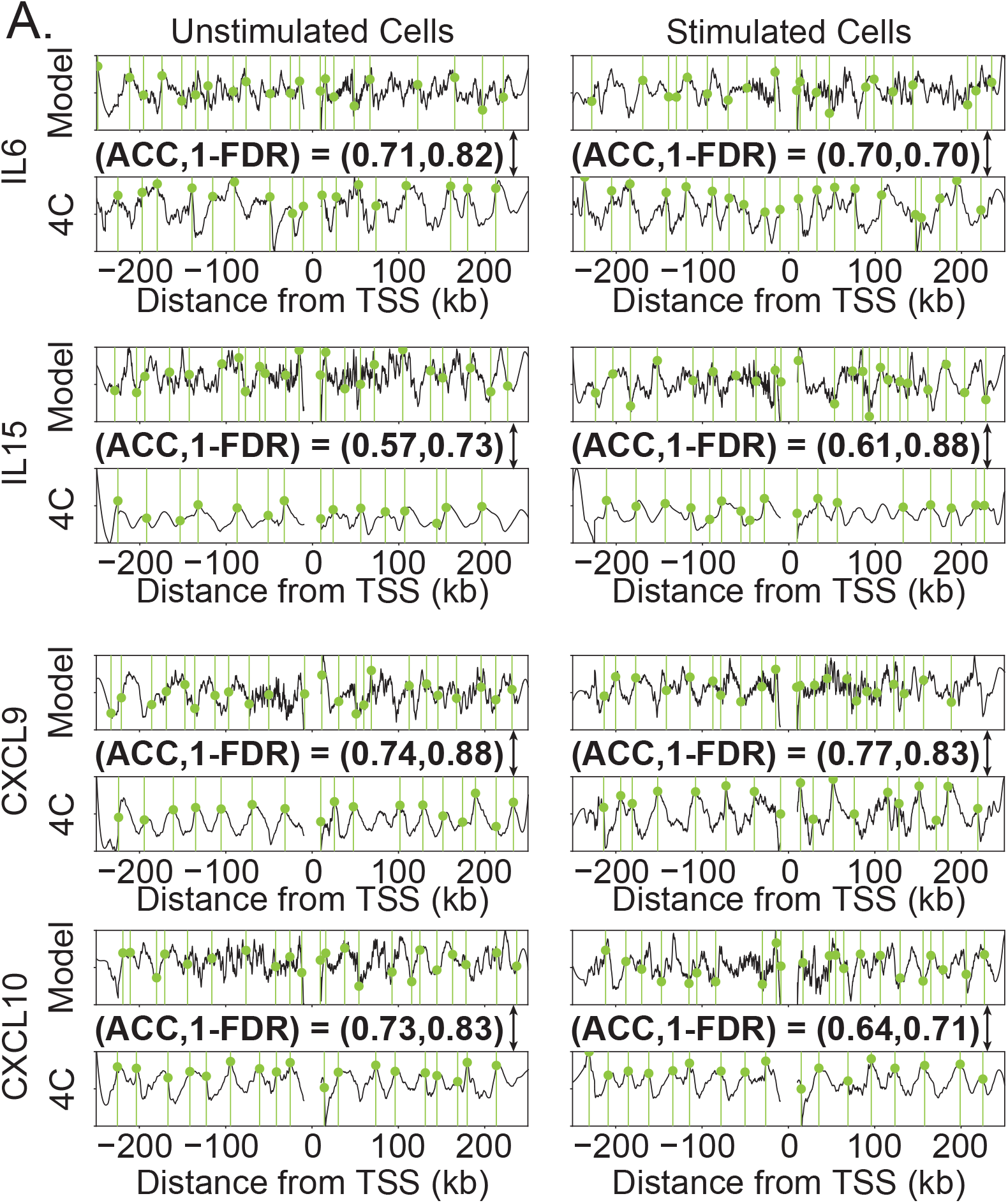
Contact signals for unstimulated, pre-LPS human macrophage (left) and post-LPS 1hr stimulated macrophage (left) for IL6, IL15, CXCL9, and CXCL10. Tracks from the ATAC-EM-informed model (top) and 4C (bottom) are shown with peaks overlayed in green. Spacing of nucleosomes in low coverage regions are distributed uniformly, as described by the model in Figure 2BE.

**Figure S19:**
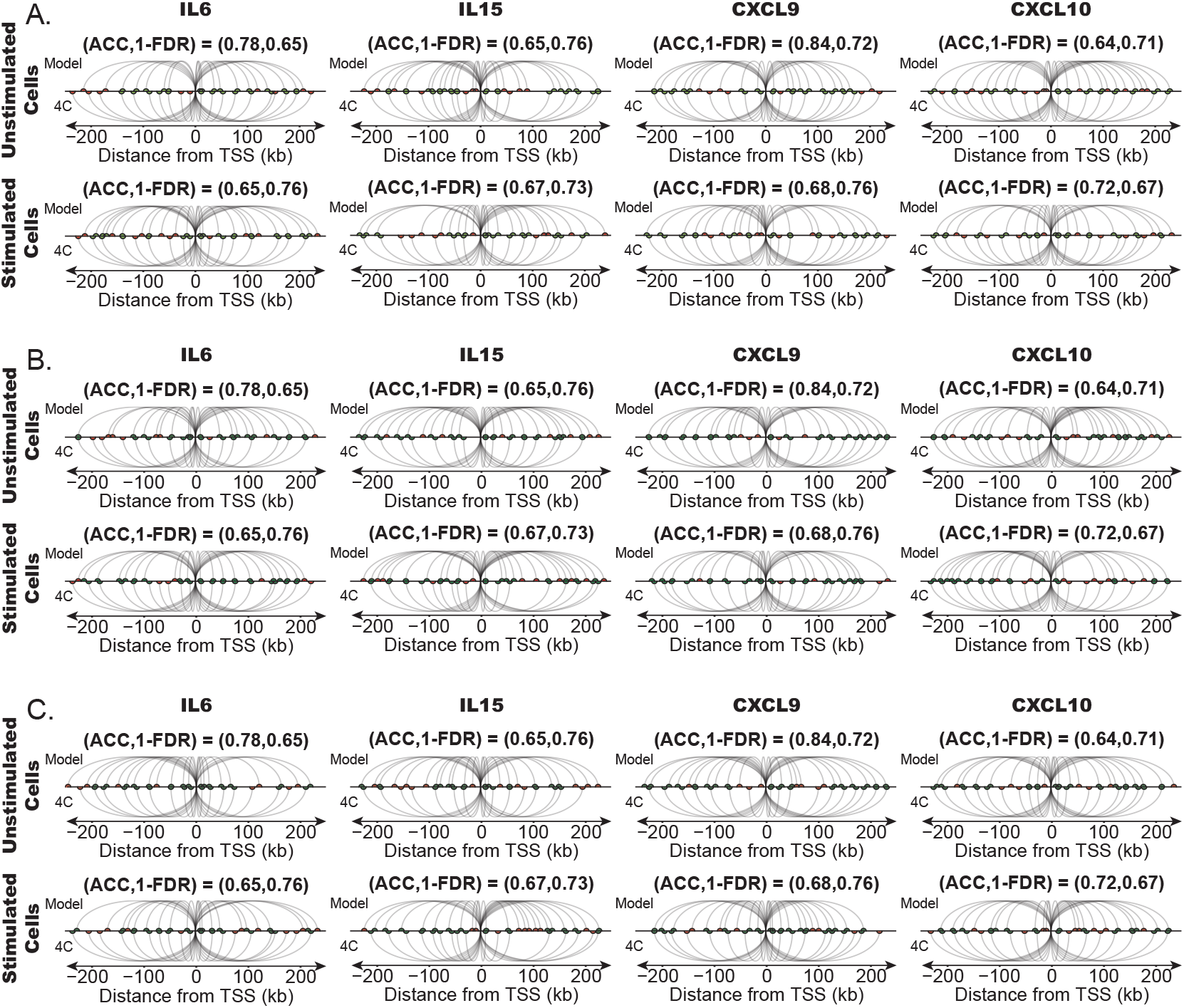
**A**. Predicted contacts for unstimulated, pre-LPS human macrophage (top) and post-LPS 1hr stimulated macrophage (bottom) for IL6, IL15, CXCL9, and CXCL10. Spacing of nucleosomes in low coverage regions are distributed exponentially, as described by the model in Figure 2CF. **B**. Predicted contacts for unstimulated, pre-LPS human macrophage (top) and post-LPS 1hr stimulated macrophage (bottom) for IL6, IL15, CXCL9, and CXCL10. Spacing of nucleosomes in low coverage regions are distributed evenly, as described by the model in Figure 2AD. **C**. Predicted contacts for unstimulated, pre-LPS human macrophage (top) and post-LPS 1hr stimulated macrophage (bottom) for IL6, IL15, CXCL9, and CXCL10. Spacing of nucleosomes in low coverage regions are distributed uniformly, as described by the model in Figure 2BE.

**Figure S20:**
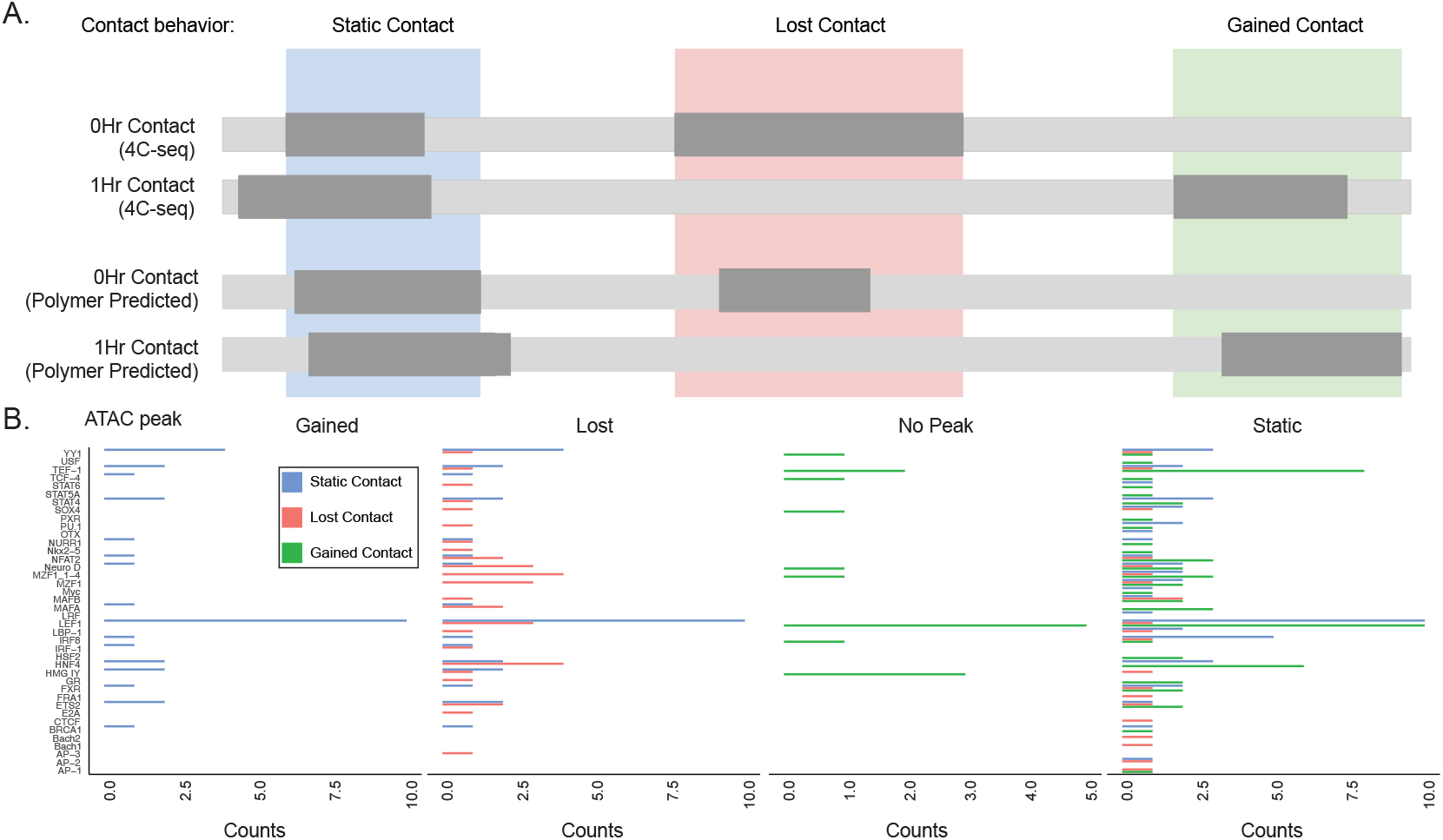
**A**. Strategy to identify polymer-mechanics-predicted and experimentally validated regions of viewpoint contacts at the CXCL9 locus. **B**. The frequency of TF binding motifs located within genomic contacts defined in (A) according to contact dynamics (gained: green; lost: red; static: blue) and divided according to the behavior of ATAC-seq peaks observed during 1hr stimulation (gained, lost, static, or no peak) overlapping with each respective contact region.

## Footnotes

1 and github.com/ajspakow/wlcstat

